# Myeloid cell networks determine reinstatement of original immune environments in recurrent ovarian cancer

**DOI:** 10.1101/2024.05.02.590528

**Authors:** Eleonora Ghisoni, Fabrizio Benedetti, Aspram Minasyan, Paula Cunnea, Alizée J. Grimm, Noémie Fahr, Mathieu Desbuisson, Charlotte Capt, Nicolas Rayroux, Doga C. Gulhan, Julien Dagher, David Barras, Matteo Morotti, Juan A. Marín-Jiménez, Flavia De Carlo, Bovannak Stewen Chap, Giulia Spagnol, Mapi Fleury, Katerina Fortis, Julien Dorier, Stephanie Tissot, Sylvie Rusakiewicz, Humberto J. Ferreira, Michal Bassani-Stenberg, Elizabeth M. Swisher, Lana Kandalft, Spyridon A. Mastroyannis, Kathleen T. Montone, Daniel Powell, Mikaël J. Pittet, Janos L. Tanyi, George Coukos, Christina Fotopoulou, Jose R. Conejo-Garcia, Denarda Dangaj Laniti

## Abstract

Immunotherapy has produced disappointing results in recurrent ovarian cancer (OC). However, the prognostic value of tumour-infiltrating lymphocytes (TILs) is largely based on the analysis of treatment-naive tumours. To understand the immunobiology of recurrent cancers, and their evolution, we profiled 170 patient-matched primary-recurrent OC samples from 69 patients of two independent cohorts. By capturing heterogeneous TIL distributions, we identified four immune phenotypes associated with differential prognosis, TILs states and TILs:myeloid networks, which dictate malignant evolution after chemotherapy and recurrence. Notably, recurrent tumours recapitulate the immunogenic patterns of original cancers. Mirroring inflamed human OC, preclinical recurrent *Brca1*^mut^ tumours maintained activated TILs:dendritic cells (DCs) niches and immunostimulatory tumour-associated macrophages (TAMs). Conversely, recurrent *Brca1*^wt^ tumours displayed loss of TILs:DCs niches and accumulated immunosuppressive myeloid networks featuring *Trem2/ApoE*^high^ TAMs and *Nduf4l2*^high^/*Galectin3*^high^ malignant states. Our study highlights that persistent immunogenicity in recurrent OC is governed by the crosstalk between dissimilar myeloid cells and TILs, which is BRCA-dependent.

## Introduction

Ovarian cancer (OC) is the leading cause of death from gynecological malignancies^1^. Despite optimal front-line treatment (cytoreductive surgery and platinum-based chemotherapy, CTX), most women with advanced-stage disease will ultimately relapse. Life expectancy for platinum-resistant patients does not exceed one year and new treatment options are urgently needed to increase response rate and survival^2^.

About half of the patients with OC contain tumour-infiltrating lymphocytes (TILs) within tumour islets, and the presence of intra-epithelial (ie)TILs in primary tumours correlates with overall survival (OS)^3–5^. Moreover, tumours which harbor homologous recombination deficiency (HRD) have a higher neo-antigen load, more TILs and up-regulation of programmed cell death protein-1 and its ligand (PD1/PD-L1) immune axis^6,7^. Despite being considered as a potential therapeutic option for OC, immune-checkpoint inhibitors (ICIs) have fallen short of expectations with no agent approved so far^8,9^. Additionally, potential biomarkers like PD-L1 expression, tumour mutational burden and TILs have yet to be proven predictive for patient selection^10^. Although the presence and state of TILs in OC have been extensively explored in previous studies^11–14^, recent research has shed light on the involvement of additional immune cell types in sculpting the tumour microenvironment (TME) of OC^15–18^. However, a comprehensive understanding of the temporal evolution of immune cell infiltration and its spatial organization in the recurrent disease setting is still lacking^13^.

Disease progression after standard-of-care therapy can lead to immune-exclusion and therapeutic failure^12,19–22^. Considering the poor outcomes of ICIs/CTX combination in the most recent trials^23–25^ it is debatable whether CTX and ICIs can effectively collaborate to promote tumour control in OC, despite quasi-universal consensus about the immunogenicity of this disease.

In this study, we employed digital pathology to capture the heterogeneity of CD8^+^ T cell infiltration among 170 OC samples and we defined four distinct tumour immune phenotypes associated with different clinical outcome and characterized by differential TILs states and TILs:myeloid networks. We demonstrated that TILs:dendritic cells (DCs) cellular crosstalk are significantly enriched at baseline and recurrence only in purely CD8^+^-inflamed tumours which carry mutations for BRCA/HRD genes and could represent ideal candidates for ICIs. Conversely, mixed-inflamed or excluded tumors harbor the highest presence of macrophages and are dominated by myeloid-cell interactions which increased further at recurrence. To mimic clinical standard-of-care, we established primary and recurrent syngeneic murine *Brca1* isogenic OC models and characterized their malignant and TME states by single-cell RNA sequencing (scRNAseq). Phenocopying inflamed human OC, *Brca1^mut^* tumours maintained activated TILs:DCs niches and further increased the infiltration of immunostimulatory tumour-associated macrophages (TAMs) at recurrence. Conversely, *Brca1^wt^* tumours displayed concomitant loss of TILs and DCs upon CTX, alike human recurrent CD8^+^-excluded OC. Instead, recurrent *Brca1^wt^* tumours were characterized by suppressive malignant states with *Nduf4l2* and *Galectin3* overexpression and dominated by immunosuppressive myeloid-cell networks including *Trem2/ApoE*^high^ TAMs and led by TGF-β-driven malignant-TAMs interactions thus resulting in a T cell-excluding phenotype.

Our study underlines that myeloid-cell networks dictate the course of immunogenic evolution of BRCA1 and HRP tumours and provides new biomarkers to improve patient selection and immunotherapy outcomes.

## Results

### Intra-tumoural heterogeneity of TILs infiltration in OC reveals four CD8^+^ immune phenotypes associated with differential prognosis

We analyzed 170 OC samples from two independent cohorts with matching treatment-näive and recurrent tumors (**Figure 1A and Table S1**). Given the absence of standardized methods for CD8^+^ T cell quantification by multiplex immunofluorescence (mIF), we built an algorithm able to capture the heterogeneity of CD8^+^ T-cell densities and their spatial distribution in whole-tissue slides. We converted the established mean of five intra-epithelial (ie)CD8^+^ T cells per high-power field captured by standardized IHC^3^ to a mean of 21 ieCD8^+^ T cells/mm^2^ quantified by mIF on whole FFPE slides (**Figure 1B**). We tested the strength of this new CD8^+^ T-cell density cut-off to discriminate OS in primary tumours from two clinical cohorts (IMCOL and UPENN, **Table S1**). Patients with tumours infiltrated by a mean of >21 ieCD8^+^ T cells/mm^2^ had significantly longer OS than those with <21 ieCD8^+^ T cells/mm^2^ (median OS 40.5 versus 27 months respectively, p=0.018) (**Figure 1C**) even after adjusting for optimal residual disease (R=0) at first surgery (**Figure S1A**).

**Figure 1:**
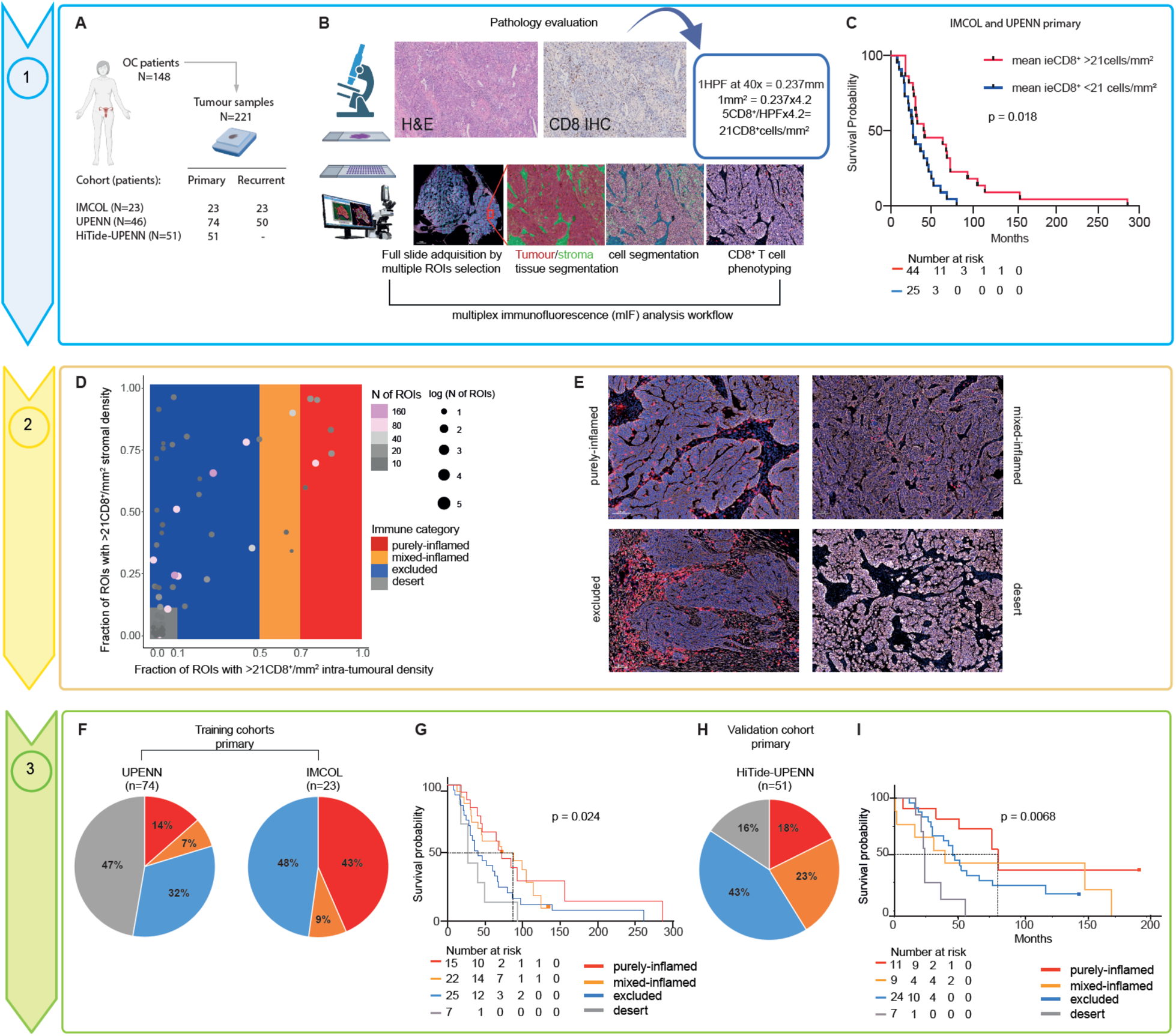
Multiplexed immunofluorescence imaging reveals four different CD8^+^-based OC immune phenotypes categories which correlate with clinical outcome. **(A)** Schematic representation of our primary-recurrent OC samples collection (N=221) coming from three different cohorts (IMCOL, UPENN and HiTide-UPENN). **(B)** Workflow of FFPE tissues analysis from pathology evaluation on hematoxylin-eosin (H&E) slides to mIF imaging. All FFPE samples (n=221, primary and recurrent OC from the three different cohort collections) were stained with mIF panel including marker for CD8^+^ T cells. Tumour and stroma regions were annotated based on CK^+^ expression and CD8^+^ T cell phenotype was assigned (Methods). A mathematic conversion of the established mean of five intra-epithelial (ie)CD8^+^ per high-power field (HPF) captured by IHC^3^ to a mean of 21 ieCD8^+^ cells/mm^2^ quantified by mIF was applied. **(C)** Kaplan-Meier curve of overall survival (OS) in the IMCOL and UPENN cohort (primary samples only) according to the new mean mIF cut-off of 21CD8^+^/mm^2^ (p value assessed using Log-rank test). **(D)** Schematic representation of our CD8^+^-based immune phenotype algorithm according to the fractions of inflamed ROIs in tumour (X axis) and stroma (Y axis) (Methods). **(E)** Representative mIF images of the four immune phenotypes, scale bar 100 um. **(F)** Pie charts representing the percentage of the four immune categories in all primary tumors of our training cohorts (UPENN and IMCOL). **(G)** Kaplan-Meier curve of OS in IMCOL-UPENN cohorts according to immune phenotypes (p value assessed using Log-rank test). **(H)** Pie chart representing the percentage of the four immune-categories of all primary tumors from the validation cohort HiTide-UPENN. **(I)** Kaplan-Meier curve of OS in the HiTide-UPENN cohort according to immune phenotypes (p value assessed using Log-rank test).

While our new mIF-based cut-off separates long-term survivors, it still represents a mean density of CD8^+^TILs and therefore ignores the observed heterogeneity across surgical specimens. Thus, we sub-segmented tissues in equal sub-regions (or regions of interest, ROIs) to cover the entire FFPE slide. We annotated tumour and stroma regions within each ROI based on pan-cytokeratin (CK^+^) expression and applied our new CD8^+^ T-cell density cut-off (21 CD8^+^ cells/mm^2^) to each ROI (**Figures S1B and S1C).**

We identified four different immune phenotypes in treatment-naïve OC: purely-inflamed, mixed-inflamed, excluded and desert tumours according to the percentage of ROIs exhibiting >21 CD8^+^ cells/mm^2^ in the intra-tumoural or stromal compartment (**Figures 1D, 1E** and Methods).

We interrogated the prevalence of these four immune phenotypes in the primary tissues of our two training cohorts (**Figure 1F**) and observed differences which could be explained by characteristics such as stage, optimal debulking rates and platinum-free interval (**Figure S1D and Table S1**). Importantly, our immune phenotype classifier significantly correlated with clinical outcome, as patients with purely and mixed-inflamed phenotypes had a statistically significant longer OS compared to those with excluded and desert tumours (median OS for purely- and mixed-inflamed was 72 and 80 months, respectively, versus 41 and 27 months for exluded and desert tumours, respectively, p=0.024) (**Figure 1G**). To validate these results, we applied our immune classifier to a third, independent OC collection (HiTide-UPENN, **Figure 1H and Table S2**). Consistently, long-term survivors were those with purely and mixed inflamed tumours (median OS for purely- and mixed-inflamed was 76 and 47 months, respectively, versus 49 and 22 months for excluded and desert, respectively, p=0.0068) (**Figure 1I**). Finally, we demonstrated that patients with purely and mixed-inflamed tumours retained better OS both in the optimally and not-optimally cytoreducted subgroups (**Figures S1E and S1F**).

As expected, immune inflammation was partly associated with HRD status as BRCA/HRD OC were more enriched in purely inflamed tumours while excluded and desert immune phenotypes were more abundant in HRP OCs (**Figures S1G, S1H and Methods**).

These results highlight that not only CD8^+^ TILs density but also their spatial distribution should be considered to classify tumour immune phenotypes more accurately in OC. Our tumour immune classifier could capture the continuum of CD8^+^ infiltration and correlate this with patients’ survival and HRD status.

### OC immune phenotypes are characterized by distinct TILs and myeloid cell states

The significant clinical association of CD8^+^ immune phenotypes with survival prompted us to investigate deeper their TILs and TME states and potentially explain how CD8^+^ T cells number and distribution in OC tissues are regulated. To do so, we focused on the HiTide-UPENN cohort (**Table S2**) for which both FFPE and snap frozen material was available (**Figure 2A**). Through mIF staining (**Figure 2B and Methods**), we observed a significant enrichment in PD1^+^CD8^+^ T cells in purely-inflamed tumours, but no differences of CD103^+^CD8^+^or GzB^+^CD8^+^ T cell proportions were observed across the four immune phenotypes (**Figures 2C, S2A and S2B)**.

**Figure 2:**
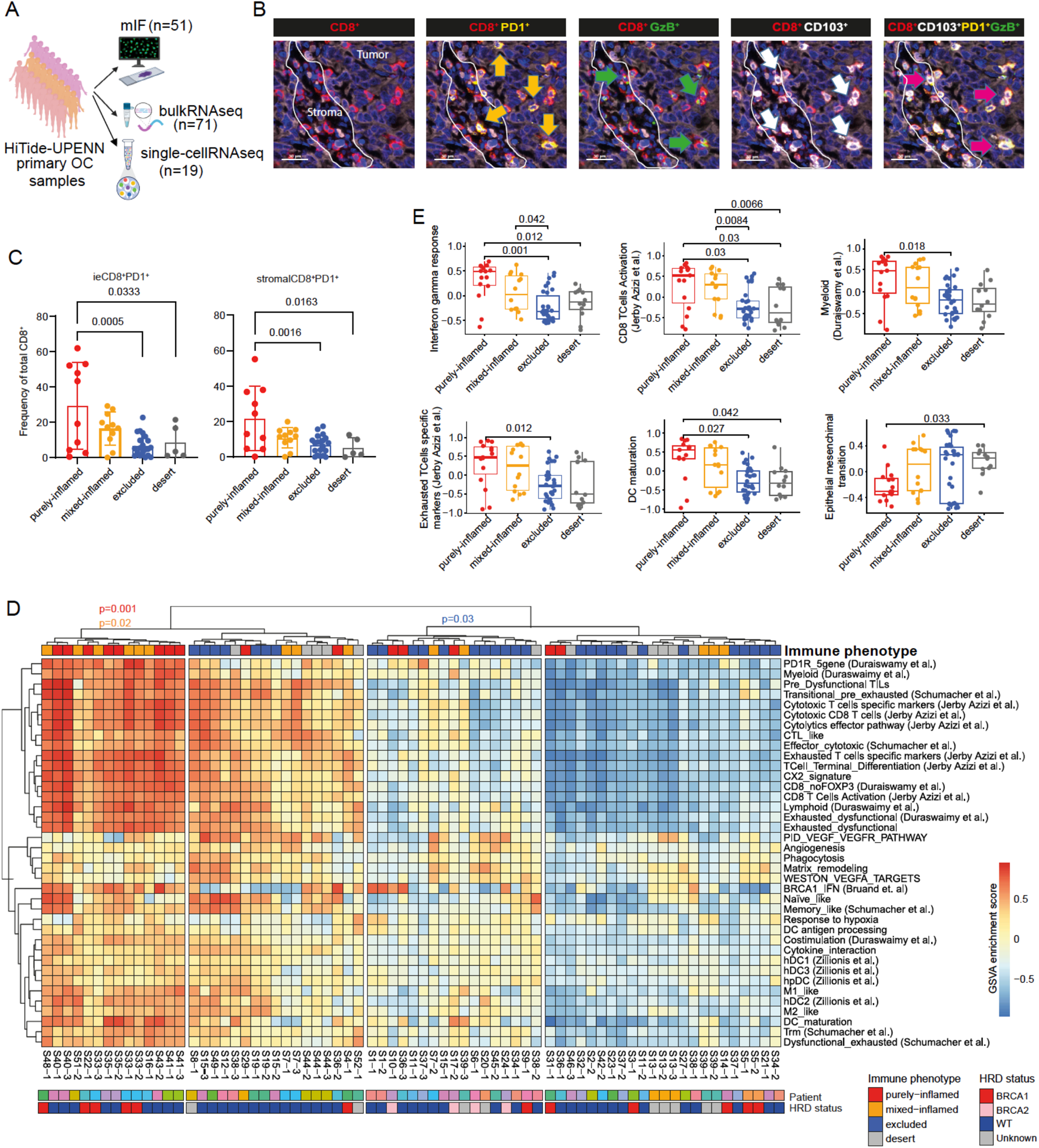
OC immune phenotypes are characterized by distinct TILs and TME states. **(A)** Schematic representation of the HiTide-UPENN primary OC cohort (n=51 patients) for which both FFPE tissue (n=51) for mIF imaging, snap frozen material (n=71) for bulk RNA and single-cell RNAsequencing (n=19) were available. **(B)** Example of the mIF panel with deconvoluted images for each marker: arrows indicate the TILs subset of interest as labelled in the upper part. **(C)** Boxplots showing proportions (%) of T cell subset of interest out of total CD8^+^ T cell profiled by mIF according to immune phenotypes and split by tumour and stromal areas (p-value assessed using unpaired, two-tailed Wilcoxon-rank test). **(D)** Heatmap of curated immune pathways (Methods and Table S2) showing different clustering among the four immune phenotypes; p-value assessed using the hypergeometric test and color-coded according to the respective immune phenotype enrichment. **(E)** Boxplots showing selected significant differential pathways among the four immune categories (extracted from panel d); error bars show the mean ± SD, only statistically significant differences shown, p-value assessed using unpaired, two-tailed Wilcoxon-rank test, adjusted for Bonferroni correction.

To gain more insight on the molecular networks characterizing each immune phenotype, we interrogated bulk RNAseq data of independent tissue sites from the same patients as above. For sixteen cases we could interrogate more than one adjacent tissue area (for example S33-1 to 3). Unsupervised clustering based on Hallmarks Reactome signatures (**Figure S2C**) revealed that the majority of purely and mixed inflamed tissues segregated together (p=0.04) and exhibited higher levels of inflammatory signatures including interferon alpha and gamma response signaling. Indeed, unsupervised clustering based on a collection of published gene signatures capturing more in depth T and myeloid cell activation^16,26–29^ (**Methods** and **Table S2)** revealed higher segregation of inflamed tissues (p=0.001 for purely-inflamed and p=0.02 for mixed-inflamed, respectively) and separated them from those assigned as excluded/desert based on corresponding mIF regions (**Figure 2D**). In a few cases (for example S36), where multiple adjacent tissues were interrogated, we observed a discrepancy in their clustering which is attributed to intrinsic intratumoural heterogeneity (ITH) often observed in OC^12,21,30^ (**Figure 2D**). Comparative bulk RNAseq analysis revealed that purely and mixed-inflamed tissues exhibited an increase in numerous T cell activation and exhaustion signatures^26–28^ (**Figures 2D and 2E**). Interestingly, inflamed tumours also exhibited higher levels of myeloid cell-related antigen presentation signatures, DC maturation^31^ and M1-macrophages^32^ (**Figure 2E**), complement and cytokine signalling^33^ denoting that TILs:myeloid cell crosstalk, crucial for T-cell engraftment and T cell costimulation^16^, were prevalent in this immune phenotype. Finally, we observed higher expression of signatures related to cancer progression and resistance to therapy in excluded and desert samples, including epithelial-mesenchymal transition (EMT), matrix remodeling, WNT-beta-catenin signaling and angiogenesis-related signatures (**Figure2E and S2D**) as previously described^34,35^. These associations could explain the observed exclusion of CD8^+^ T cell from tumour islets and the worse prognosis linked to these immune phenotypes.

In conclusion, we show that OC immune phenotypes exhibit not only different density and spatial CD8^+^ T cells infiltration but also phenotypically divergent TILs and myeloid cell states which could contribute to the observed differential clinical outcomes.

### TILs:myeloid crosstalk varies vastly across OC CD8^+^ immune phenotypes

A mounting body of evidence indicates that the state of terminally exhausted ieCD8^+^ TILs may vary depending on their cellular interactions with myeloid cells^16,36,37^.

To understand if the enrichment of antigen experienced/exhausted CD8^+^ TILs in inflamed samples is indeed sustained by the presence of intratumoural myeloid cell states, we analyzed treatment-naïve specimens from our two training cohorts (**Figure 3A and Table S1**) by mIF (**Figure 3B**). We found higher CD11c^+^ density in purely-inflamed samples compared to mixed or excluded cases in the tumour compartment but equal or even higher CD11c^+^ densities (with or without PD-L1^+^) in tumour and stroma regions of desert cases (**Figure 3C**), suggesting that differential subsets of CD11c^+^ myeloid cells must reside in inflamed and desert cases which cannot be captured merely by one marker. When analyzing the total infiltration of CD68^+^ tumour associated macrophages (TAMs) in the UPENN cohort, we observed that they were quasi-universally present in OC cases with the exception of five purely-inflamed samples where they were completely absent (**Fig.3d**).

**Figure 3:**
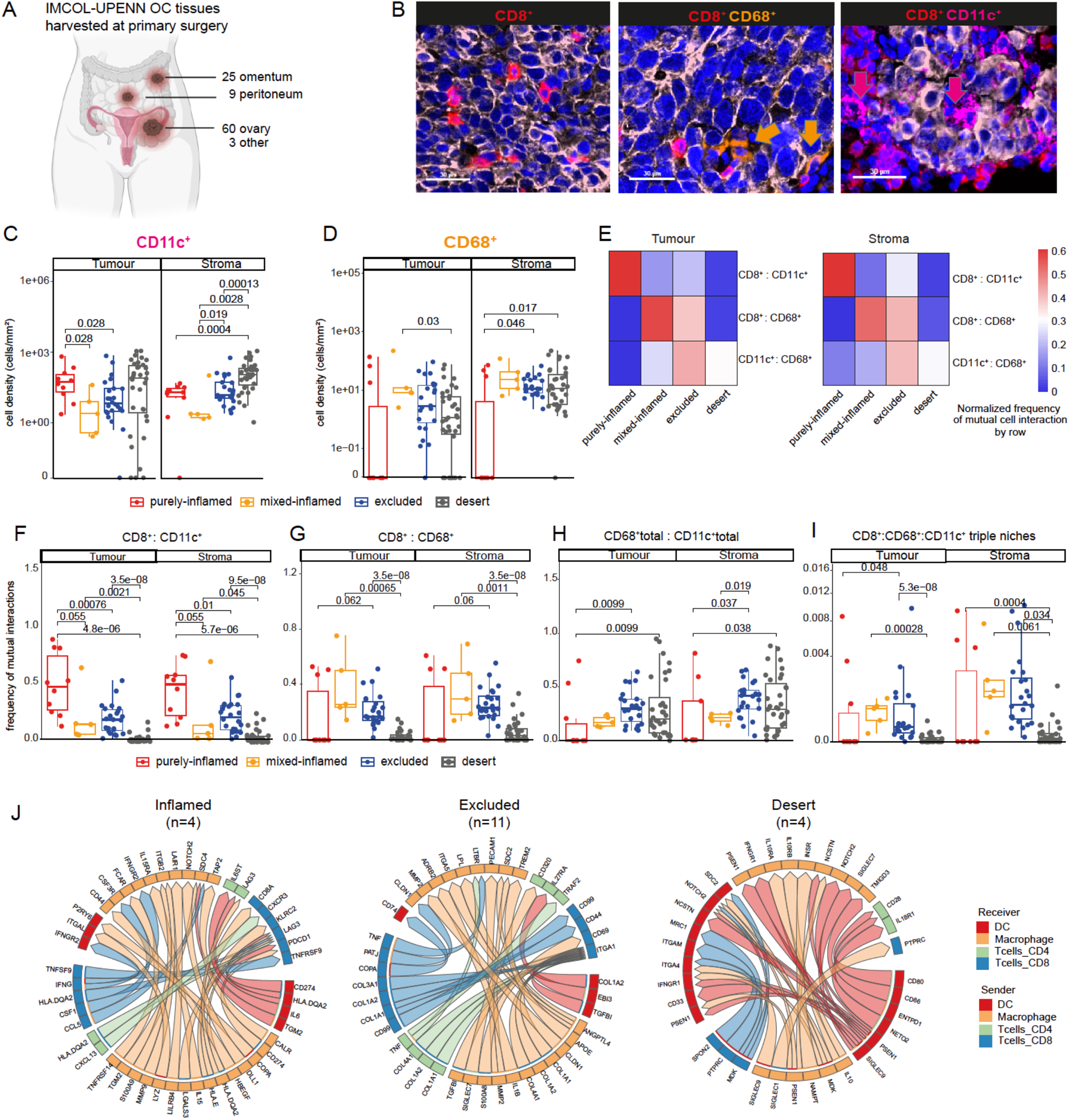
TILs: myeloid crosstalk varies vastly across OC CD8^+^ immune phenotypes. **(A)** Schematic representation of the tissue site location for samples harvested at primary surgery (treatment-naïve primary tumours for the UPENN and IMCOL, cohorts merged). **(B)** Example of the mIF panel with deconvoluted images for each marker: the immune population of interest is indicated by arrows and color-coded as labelled in the upper part: CD8 in red, CD68 in orange, CD11c in purple. **(C-D)** Boxpolts showing cell density (cell/mm^2^) profiled by mIF for CD11c^+^ and CD68^+^ according to immune phenotypes and split by tumour and stromal areas in the UPENN cohort; error bars show the mean ± SD, only statistically significant differences shown, p-value assessed using unpaired, two-tailed Wilcoxon-rank test. **(E)** Heatmaps showing the normalized frequency of mutual cell interaction at a 20um neighboring radii in the tumour and stroma compartment in the UPENN cohort (primary tumours). Immune cell population interaction of interest in lines and immune categories as columns. Color-code scale bar showing the normalized frequency by row. **(F-I)** Boxplots displaying frequency of mutual interaction between the indicated cell types according to immune phenotypes and split by tumour and stromal areas; error bars show the mean ± SD, only statistically significant differences shown, unadjusted p-value assessed using unpaired, two-tailed Wilcoxon-rank test. **(J)** Circos plot of MultiNicheNet analysis (from the HiTide-UPENN cohort subset, n=19) displaying best 30 ligand interactions per immune category for T cell, DCs and macrophages.

To gain more insight into differential TME architectures, we interrogated cellular crosstalk between CD8 ^+^ TILs and myeloid populations. Our work and others have recently shown the existence of intratumoural niches where critical T cell–DC interactions occur^16,38,39^. We derived a mutual cell-to-cell interaction neighborhood algorithm (**Figures 3E, S3A and Methods)** which calculates the normalized frequency of cell-to-cell interactions among two^37^ or more cell types (**Methods**).

We showed that purely inflamed samples exhibited significantly higher CD8^+^:CD11c^+^ interactions in both the tumour and stromal compartment compared to all other immune phenotypes (**Figure 3F**). Instead, mixed inflamed and excluded tumours harbored higher CD8^+^:CD68^+^interactions (**Figure 3G**). Although cell frequencies and their mutual interaction are interdependent by design, we found that CD8^+^:CD11c^+^ cells interaction separated purely-inflamed samples from other immune phenotypes significantly better than their respective minimal cell frequency (**Figure 3C**). Of note, we also observed that excluded and desert tumours exhibited increased CD11c^+^:CD68^+^ (or homotypic myeloid) interactions (**Figure 3H**) and wondered if those myeloid niches could also harbor CD8^+^ TILs. We thus computed the occurrence of triplet niches *in situ* (**Methods**). Our results showed that mixed-inflamed and excluded samples had higher levels of triplets populated by CD8^+^:CD11c^+^:CD68^+^ in the tumour and even more in the stroma (**Figures 3I and S3B**). This led us to hypothesize that some TAM states could interfere with productive CD8^+^:CD11c^+^ interactions thus impairing T cell co-stimulation.

When extending our analyses to include T cells with PD1 or myeloid cells with PD-L1 expression, we confirmed an enrichment of CD8^+^PD1^+^ cells interacting with CD11c^+^ cells expressing or not PD-L1 in purely-inflamed, suggesting the relevance of PD1/PDL1 axis in this OC subgroup (**Figure S3C**). On the contrary, CD11c^+^:CD68^+^PD-L1^+^ niches were absent in purely inflamed cases (**Figure S3D)** and, importantly, patients with primary tumours enriched in myeloid niches marked by PD-L1 expression had a significantly worse progression-free survival (PFS) compared to those with low CD11c^+^:CD68^+^PD-L1^+^ niches (**Figures S3E and S3F**).

To further dissect the specific immune interactions mediating the above niches we used MultiNicheNet^40^ analysis on a subset of HiTide-UPENN primary samples (N=19) for which single-cell data were collected. We showed that in inflamed samples T:myeloid cell interactions were sustained by antigen presentation (*HLA* molecules) and stimulatory signaling (*CD8A*, *IFNG),* myeloid cell recruitment *(CSF1,CCL5)* as well as by known checkpoint ligands (*PD1*, *LAG3*). Excluded tumours exhibited homotypic myeloid and T:myeloid networks dominated by collagen molecules interacting with *ITGA1*, Claudins (*CLDN1*) and *TGFB,* which are known to enhance migration and invasion in OC cells^41^ and the inhibitory *TREM2*/*APOE* axis. Finally, desert samples displayed inhibitory networks mediated by *SPON2*, *MRC1*, *PSEN1*, *NOTCH*, *ITGAM* and *SIGLEC*^42,43^ interactions (**Figure 3J and Table S3**).

These results highlight the differential CD8^+^:myeloid crosstalk established among the four immune phenotypes and shed light in their molecular interactome. The subset of purely-inflamed OC is selectively enriched for CD8^+^:CD11c^+^ niches recently shown to be essential for response to ICIs^16,39,44^ and adoptive T cell therapy^37^. They also suggest that the type of myeloid cells infiltrating tumours could be regulating T cell distribution in the TME^45^.

### HRD status and TILs:myeloid crosstalk define OC immune phenotype evolution and architecture at recurrence

We then sought to decipher the evolution of immune phenotypes upon standard-of-care CTX and recurrence taking advantage of our patient-matched tumour tissues harvested at secondary cytoreductive surgery (**Figure 4A and TableS1**).

**Figure 4:**
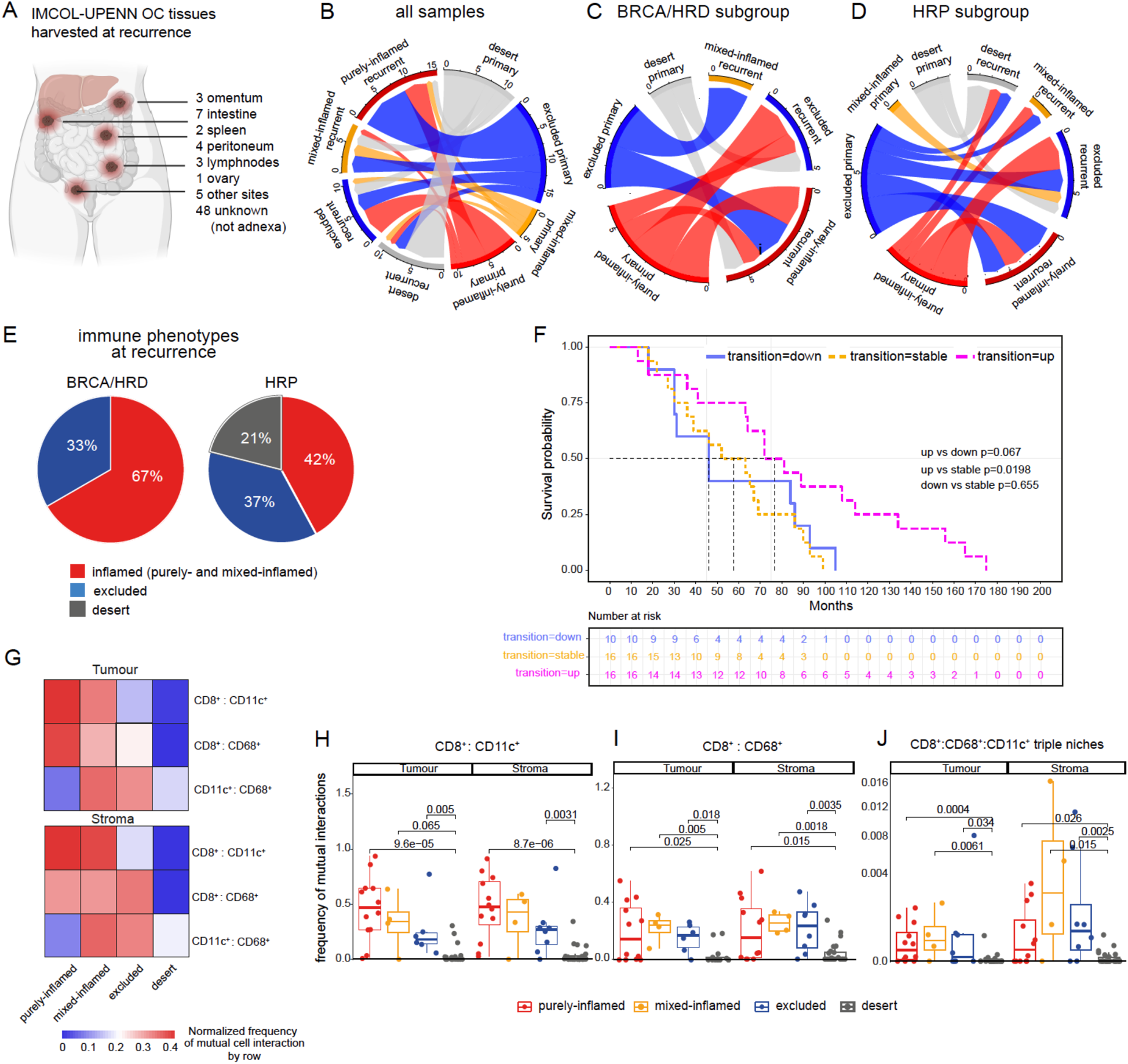
Myeloid crosstalk are upregulated at recurrence defining the evolution of OC TME architecture together with HRD status. **(A)** Schematic representation of the tissue site location for samples harvested at recurrence (UPENN and IMCOL recurrent tumours, cohorts merged). **(B-D)** Circos plots showing the evolution of immune phenotypes for patient-matched samples (UPENN and IMCOL cohorts merged) and in the BRCA/HRD and HRP subgroups separately. **(E)** Pie chart representing the percentage of the immune phenotypes at recurrence in the BRCA/HRD and HRP subgroups. **(F)** Kaplan-Meier curves of OS according to immune phenotype evolution at recurrence (p value assessed using Log-rank test). **(G)** Heatmaps showing the normalized frequency of mutual interaction between cell types at a 20um neighboring radii in the tumour and stroma compartment at recurrence for UPENN cohort. Immune cell population interaction of interest in lines and immune categories as columns. Color-code scale bar showing the normalized frequency by each row. **(H-J)** Boxplots displaying frequency of mutual interaction between the indicated cell types according to immune phenotypes and split by tumour or stromal or areas; error bars show the mean ± SD, only statistically significant differences shown, unadjusted p-value assessed using unpaired, two-tailed Wilcoxon-rank test.

We applied our immune classification in recurrent OC tissues and observed that the trajectory and evolution of immune phenotypes was highly dynamic (**Figure 4B**). Nevertheless, most purely-inflamed OCs retained their homogenous CD8^+^ inflammation whereas most desert carcinomas remained desert upon recurrence, suggesting that re-emerging tumours could reconstitute their CD8^+^ spatial distribution and, by extension, their TME.

Indeed, when tracking the evolution of immune phenotypes according to HRD status, we showed that whereas HRP tumours largely spread across different immune phenotypes and toward an excluded or desert phenotype, most recurrent BRCA/HRD retained or even evolved toward an inflammatory state (**Figures 4C-4E**). Importantly the evolution toward an inflamed phenotype at recurrence was associated with a benefit in OS (**Figure 4F**).

When interrogating TILs:myeloid cell neighborhoods we saw that recurrent purely-inflamed tumours retained the highest frequency of CD8^+^:CD11c^+^ interactions (**Figures 4G and 4H**). Mixed-inflamed cases were enriched in both CD8^+^:CD11c^+^ and CD8^+^:CD68^+^ interactions (**Figure 4I**). Finally, recurrent excluded tumours were mostly enriched by CD8^+^:CD68^+^ and triplet (CD8^+^:CD11c^+^:CD68^+^) niches similarly to primary OC (**Figures 4I and 4J**).

Despite the limitation of having patient-matched but not site-matched samples, our analyses revealed that immune cell dynamics are affected by disease progression, but the key TME players and interactions are rather stable at recurrence in OC immune phenotypes. Purely-inflamed tumours maintain homogenous TIL inflammation in line with the presence of CD8^+^:DC crosstalk in these tumours, while mixed inflamed and excluded OCs exhibit higher TILs:TAMs and homotypic myeloid interactions, thus suggesting a faster evolution toward an immune-resistant phenotype with rare tumor-reactive resident TILs able to to abrogate malignant progression^11,46^. Our data also suggests that HRD mutational status could potentially determine tumour immune phenotype evolution and thus warrants the further investigation following below.

### Temporal heterogeneity of T:myeloid cell inflammation in recurrent mouse OC models

To further disentangle the molecular mechanisms underlying TIL infiltration and TME orchestration we set out to build orthotopic mouse OC models with defined HRD status which resemble primary and recurrent human OCs as well as their respective TMEs. We employed the syngeneic ID8 cell lines knocked out for *Trp53* and *Brca1* genes^47,48^ (**Methods**) and further engineered to overexpress luciferase^29^. Mice were orthotopically implanted with *Trp53*^−/−^ *Brca1*^−/−^ (hereby referred as *Brca1*^mut^) or *Trp53*^−/−^ *Brca1*^+/+^ (or *Brca1*^wt^) ID8 tumour cells and treated weekly with dual CTX (carboplatin/paclitaxel) for 6 cycles, mimicking first line clinical standard-of-care. Bioluminescence abdominal quantification revealed partial or even complete tumour regression upon CTX in mice, before they all eventually recurred with *Brca1*^mut^ having a slower relapse kinetic than *Brca1*^wt^ (**Figures 5A and S4A)**.

**Figure 5:**
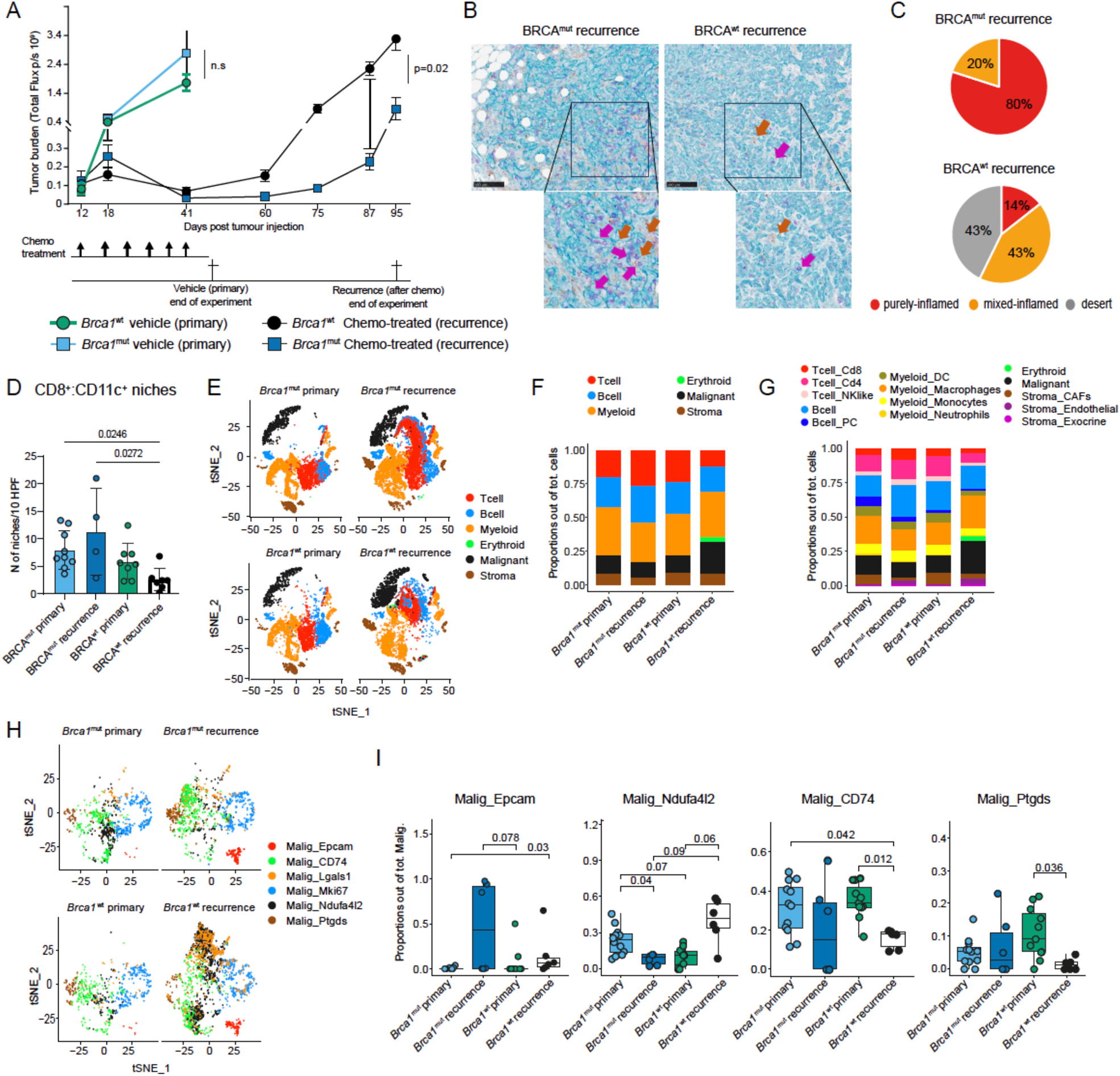
*Brca1*^wt^ tumours lose TILs:APC interactions at recurrence and express suppressive malignant states. **(A)** Tumour growth kinetics of *Tpr53^−/−^* ID8Luc *Brca1*^wt^ and *Brca1*^mut^ during treatment in the control (vehicle or primary in the figure) or chemotherapy group (recurrence) with weekly carboplatin-taxol i.p injection for 6 weeks total (n = 6-7 mice per group); p values calculated by a two-way ANOVA. **(B)** Multiplex IHC staining of *Brca1*^mut^ and *Brca1*^wt^ tumours. Arrows showing examples of CD8+ (purple) and CD11c+ (brown). Original scale bars, 100 mm. **(C)** Pie-charts displaying the different percentage of immune phenotypes at recurrence (after chemotherapy treatment) between *Brca1*^mut^ and *Brca1*^wt^ tumours. **(D)** Boxplots showing CD8^+^:CD11c^+^ niches assessed by IHC between *Brca1*^mut^ and *Brca1*^wt^ tumours at baseline and at recurrence; p-value assessed using unpaired, two-tailed Wilcoxon-rank test. **(E)** t-SNE map of single-cell transcriptomic data displaying the identified 6 major cell clusters among *Brca1*^mut^ and *Brca1*^wt^ tumours at baseline and at recurrence. **(F-G)** Bar plots displaying respectively the proportion of major cell type and subtypes (out of total cells) among *Brca1*^mut^ and *Brca1*^wt^ tumours at baseline and at recurrence. **(H)** t-SNE map of single-cell data reveals 6 major malignant clusters subtypes. **(I)** Boxplots displaying the proportion of indicated malignant subclasses between *Brca1*^mut^ and *Brca1*^wt^ tumours at baseline and at recurrence; p-value assessed using unpaired, two-tailed Wilcoxon-rank test, adjusted for Bonferroni correction.

To immune-classify and study TILs:DCs crosstalk in our primary and recurrent mouse models, we performed a triple IHC staining for CD8^+^, CD11c^+^ and panCK^+^ cells in mouse tumor tissues (**Figure 5B**). *Brca1^mut^*displayed higher levels of CD8^+^ and CD11c^+^ densities at baseline and maintained them at recurrence while strikingly *Brca1^wt^*lost both CD8^+^ TILs and CD11c^+^ (**Figure S4B**). Only *Brca1*^mut^ remained homogenously inflamed at recurrence with concomitant higher CD8^+^:CD11c^+^ niches (**Figures 5C, 5D and S4C**). However, recurrent *Brca1*^wt^ tumours evolved mainly into desert phenotypes (17% at primary vs 43% at recurrence) with a global depletion of CD8^+^:CD11c^+^ niches (**Figures 5C and S4C**). The above findings indicated that *Brca1^mut^* tumours recapitulate the low percentage of purely-inflamed cases observed in the human dataset characterized by homogenous CD8^+^ TIL infiltration and high number of TILs:APC interactions. On the contrary, *BRCA^wt^* (mainly desert at recurrence) may reflect CTX-resistant tumours where an immune-suppressive TME with loss of TILs and DCs develops upon progression^49^.

These findings prompted us to further dissect and compare the evolution of the TME in our primary-recurrent OC models by scRNAseq. We analyzed 39’752 cells distributed into 17 major clusters as displayed in our t-distributed stochastic neighbor embedding (tSNE) plots (**Figure S4D and Methods**) and identified six major cell classes (**Figures 5E, 5F and S4E**) and 14 subclasses (**Figures 5G and S4F**). We validated by scRNAseq that purely-inflamed mouse OCs exhibited higher proportions of T cells and myeloid DC cells compared to mixed-inflamed and desert samples (p=0.017 and p=0.051, respectively) while mixed and desert samples were enriched in myeloid macrophages and malignant subsets (**Figures S4G and S4H**) recapitulating our human data previously shown.

When studying the malignant and stromal compartment of our models we observed divergent TME compositions and evolution at recurrence. *Brca1*^mut^ tumours were dominated by malignant cell states such as *CD74*^high^ and *Ptgds*^high^ clusters (**Figures 5H, 5I, S4I and Table S4)** with overexpression of FGFR signaling pathways but also antigen processing/presentation genes (MHC class II [*H2-Eb1*, *H2-Ab1*, *H2-Aa*], *Cd74*, *Cd86*) **(Figures S4J and S4K)**. *Ptgds*^high^ cluster was also characterized by prostanoid and eicosanoid-associated metabolic signatures and increased specifically after CTX and tumour progression (**Figures 5H, 5I, S4J and S4K**). Recurrent *Brca1^mut^* also maintained a highly proliferating *Mki67* ^high^ tumour cell state with overexpression of genes such as *Hmgb2* and senescence-associated secretory phenotype (SASP)^50^. However, recurrent *Brca1^wt^* significantly lost the *CD74* ^high^, *Mki67* ^high^ and *Ptgds*^high^ malignant cell states and were largely repopulated by the immunosuppressive *Nduf4l2*^high^ compartment overexpressing *Lgals1*^51^ (**Figures 5H, 5I and S4K**). Reactome analysis showed that this state was enriched for expression of signatures of EMT–driven phosphatidylinositol 3-kinase (PI3K)–AKT pathway, NCAM1 and LG1-ADAM interactions and to a lesser extent, STING-mediated immune-response signatures (**Figure S4J**).

Our data on the malignant compartment evolution may explain why *Brca1^mut^*tumours remain immunogenic at recurrence and therefore maintain their TILs:DCs niches but can still evade immune destruction through FGFR^52^ and COX-driven prostanoid signaling^53^. In contrast, *Brca1^wt^* tumours are highly reshaped during tumour progression. They emerge into new immune evading and suppressive malignant states with *Nduf4l2* and *Galectin3* overexpression ^51,54,55^. These results may explain the observed loss of T cells and stimulatory APCs which indeed leads to faster progression in *Brca1^wt^* tumours as observed in end-stage human OC^13^.

To further understand how the stromal compartment accompanies these malignant-cell evolution we compared changes of endothelial and fibroblast populations at baseline and recurrence. In both tumour models, we observed a striking increase of endothelial cells after CTX, suggesting that cancer progression drives angiogenesis (**Figures S5A and S5B**). *Brca1^mut^* and *Brca1^wt^* tumours both lost proliferating (*Ki67*^high^) cancer-associated fibroblasts (CAFs). In addition, recurrent *Brca1^mut^* reshaped their CAF composition by significantly reducing the clusterin^high^ (*Clu*^high^)-CAFs cluster which modulate the adjacent TME via TGFβ signaling^56^ and by increasing the inflammatory *Gsn*^high^-CAFs cluster (**Figures 6A, S5A and Table S4**), reported to overexpress multiple pathways involving *Ptgis* (Prostaglandin I2 synthase) and complement activation through C3 and CFD^57^ (**Table S4**).

**Figure 6:**
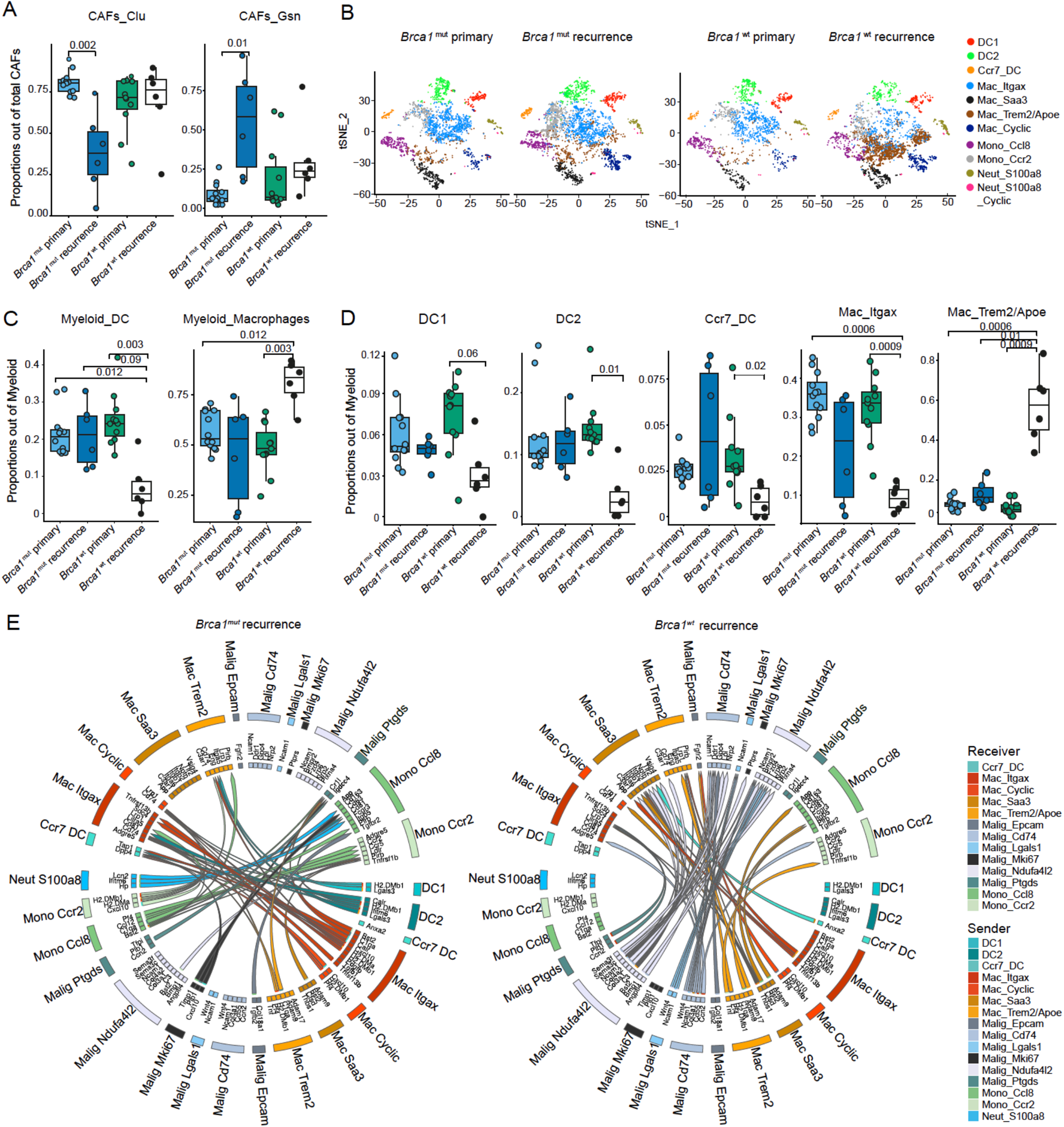
*Brca1*^wt^ upregulate immunosuppressive TAMs and CAFs at recurrence and are dominated by macrophages-malignant crosstalk. **(A)** Boxplots displaying the proportion of indicated CAFs subclasses between *Brca1*^mut^ and *Brca1*^wt^ tumours at baseline and at recurrence; p-value assessed using unpaired, two-tailed Wilcoxon-rank test, adjusted for Bonferroni correction. **(B)** t-SNE map of single-cell data reveals 11 major myeloid subtypes. **(C-D)** Boxplots displaying the proportion of indicated Myeloid classes and subclasses between *Brca1*^mut^ and *Brca1*^wt^ tumours at baseline and at recurrence; p-value assessed using unpaired, two-tailed Wilcoxon-rank test, adjusted for Bonferroni correction. **(E)** Circos plot of interactome analysis by MultiNicheNet displaying finer subclasses interaction within the top 5 cell type interaction between *Brca1*^mut^ and *Brca1*^wt^ recurrent tumours.

These results highlight highly divergent evolutions in the malignant and stromal landscape of HRD and HRP recurrent tumours and could further explain the vast heterogeneity observed in the temporal tumor immune phenotype evolution of human OC. In summary, recurrent *Brca1*^mut^ tumours retain an immunogenic malignant compartment while *Brca1*^wt^ tumours evolve into immune evasive malignant states.

### *Tgfb1, Fn1, ApoE/Trem2 and Thbs1* signaling drive immunosuppressive TAM networks in recurrent *Brca1^wt^* tumours

While we could observe a global loss of T cells in *Brca1^wt^*and their maintenance in *Brca1^mut^* tumours (**Figures 5F, 5G and S5C**), more subtle changes appeared in their composition at recurrence. The CD8 and CD4 naive-like, effector memory and exhausted TIL subsets remained unaltered (**Figure S5C**), while the CD4-resting state increased (**Figure S5D**). In addition, *BRCA1^mut^* recurrent tumours maintained a higher type-I interferon CD8 TIL state which was lost in *Brca1^wt^*and increased the frequency of *Hsp*^high^ CD8 TIL state (**Figures S5E and S5F**).

Having described a clear interplay between myeloid cells and TILs in our human dataset, we investigated the evolution of myeloid cell subtypes in our mouse models. Comparative analysis showed a drastic decrease of DCs affecting all subsets (cDC1, cDC2, CCR7^+^DC) in recurrent *Brca1^wt^* tumours (**Figures 6B-6D and S5G**). This was counterbalanced by a global increase in macrophages and specifically by *Trem2*/*ApoE* TAM state^58,59^ (**Figures 6C and 6D**) characterized among others by stress-induced senescence and TNFR1-driven NF-k beta signaling and involved in HDL metabolism^60,61^ (**Table S4**). A clear dichotomy was observed between our two models at recurrence where *Itgax*-TAMs overexpressing *Cxcl9*, *Cxcl16* and *Il1b* were completely lost at recurrence in *Brca1^wt^* tumours while they were maintained, together with all the DC subsets, in *Brca1^mu^*^t^ cancers (**Figures 6C, 6D and S5H**).

The above findings prompted us to predict the signaling networks and infer the cellular crosstalk established in the TMEs of these temporally divergent tumour immune phenotypes.

By applying MultiNicheNet^40^ we revealed highly divergent regulatory networks at recurrence between our models (**Figure S6A**). While *Brca1^mut^* displayed a myeloid:T cell network sustaining antigen-presentation and chemokine activation, a plethora of inhibitory macrophages-malignant and homotypic myeloid cell signaling and interactions dominated *Brca1^wt^* tumours with downregulation of antigen presentation. To increase resolution in the interactome, we next focused on the top five cell-type ligand-receptor interactions of each recurrent tumour model. Again, we found that DCs:B cells:CD4 T cells interactions were maintained in *Brca1^mut^* upon recurrence and lost in *Brca1^wt^* (**Figure S6B and Table S5**). These were sustained by the highly important *Cxcl9/Cxcl10*-*Cxcr3* axis for OC^33^ and provide TILs co-stimulation through CD28^16,33^. On the contrary, *Brca1^wt^* tumours were dominated by homotypic myeloid-cell crosstalk including macrophages:DCs predicted to interact predominantly through *Trem2*/*Apoe*^high^ TAMs and mediated by *Tgfb1*, *Fn1*, *App* and *Thbs1* ligands associated with EMT, invasiveness and metastatic spread^62,63^ (**Figure 6E**).

In conclusion, our findings demonstrated that *Brca1^mut^*retain their tumour immune phenotype due to immunogenic malignant and potentially immune stimulatory myeloid cell subsets. On the contrary, *Brca1^wt^*(mainly desert at recurrence) just like chemo-resistant tumours establish a TIL-excluding TME due to the loss of activated DCs. In addition, phenocopying tumour progression of immune excluded OC, they reveal upregulation of inhibitory *Trem2*/*Apoe*^high^ TAMs subsets, potentially recruited by immune evasive *Nduf4l2/Galectin3*^high^ malignant states.

## Discussion

ICIs have revolutionized the immuno-oncology field but have failed to demonstrate efficacy in OC despite the ample evidence of adaptive immunity being activated at baseline. The discrepancy could rely on the poor understanding of the mechanisms that regulate the temporal evolution of TILs and myeloid cell networks with disease progression, known to be relevant for clinical response.

To capture and quantify the breadth of TIL infiltration in OC, we applied digital pathology mIF analysis on whole tissue slides and built a tumour immune phenotype predicting algorithm which systematically classified 221 OC specimens (from 2 training and 1 validation cohort) in four CD8^+^-based immune-categories. Importantly patients with purely-inflamed OC showed better OS and carried the highest levels of CD8^+^PD1^+^ antigen-experienced/exhausted TILs. In addition, these tissues were characterized by increased interferon gamma and alpha activation and accompanied by activated myeloid signatures. Spatial neighborhood analysis further revealed that this small OC subset harbored intratumoural TILs:DCs niches important for response to ICIs in ovarian but also to adoptive T cell therapy in melanoma^37^. Altogether our results suggest that the small subgroup of purely-inflamed OC identified by our algorithm could represent the ideal candidates for immunotherapy trials.

Conversely, mixed-inflamed, excluded and desert tumours were enriched in TILs:TAMs or myeloid homotypic myeloid interactions which were associated with worse outcomes implying that TAMs could interfere with stimulatory and proficient T cell:DC interactions or exert a “trapping effect” of either pair population.

Upon CTX pressure and recurrence, about half of OC preserved or restored their tumour immune phenotype and TILs:DCs crosstalk and those were more frequently enriched in mutations of the homologous recombination repair pathway. Notably, tumours which amplified their TIL content at recurrence had an improved survival.

However, the other half of OC stochastically evolved in any tumour immune phenotype. Recurrent mixed-inflamed, excluded and desert tumors increased the levels of TILs:TAMs or homotypic TAMs crosstalk, highlighting the important role of this myeloid cell evolution at tumour progression, in line with multiple studies in ovarian and other cancer types^64,65^.

Systematically classifying the tumour immune phenotype and predicting the factors that stabilize or enrich T cells or those which exclude them from the TME at recurrence holds value for the appropriate choice of therapeutic agents upon first line treatment.

Our findings suggest that both tumour-intrinsic and extrinsic mechanisms play a defining role in the dynamics of TILs infiltration patterns at recurrence. To mechanistically disentangle TME at tumour progression, we translated the clinical standard-of-care treatment of OC in preclinical syngeneic mouse *Brca1* isogenic ovarian cancer models, and comparatively characterized the evolution of their malignant and TME states. Our results revealed strikingly different temporal evolutions between *Brca1^mut^*and *Brca1^wt^* recurrent tumours, underscoring the need for differential treatment strategies for these subgroups.

Phenocopying inflamed human OC, *Brca1^mut^* tumours maintained activated TILs:DCs niches at recurrence and further increased the infiltration of immunostimulatory TAMs. This was likely enabled by immunogenic tumour cell states with increased antigen presentation and inflammatory CAFs. However, they could still evade T cell-mediated destruction likely due to the upregulation of PGE2-producing signaling pathways known to restrict the expansion of antigen-experienced TILs and downstream destruction of IL-2 signaling and metabolic fitness impairment^66,67^. Our data also identify *Brca1^mut^*recurrent OC as potential candidate for blocking the PGE2-EP2/EP4 axis to unleash the functionality of the TILs:DCs niches.

In contrast, *Brca1^wt^* tumours displayed concomitant loss of TILs and DCs upon CTX, alike human recurrent CD8^+^-excluded OC. Instead, they were highly infiltrated by TAMs reprogrammed to overexpress the *Trem2*/*ApoE* axis involved in HDL metabolism. These was likely driven by emerging suppressive and highly metabolic malignant states with *Nduf4l2* and *Galectin3* overexpression^51,54^ and characterized by signatures of the EMT-PI3K–AKT pathway, NCAM1 and LG1-ADAM interactions^68^. *Brca1^wt^*tumors were also dominated by immunosuppressive homotypic myeloid-cell and malignant:TAMs crosstalk mediated by *Tgfb1*, Fn1, *App* and *Thbs1* signaling which govern an immunosuppressive TME^62^ and are associated with resistance to ICIs^69^.

Our findings provide important mechanistic insights about the complex spatial and temporal evolution of the TME of OC and provide new targets for differential treatment approaches according to BRCA/HRD status. Furthermore, they underscore that both the mutational status and CD8^+^ tumour immune phenotype should be taken into consideration as combined biomarkers when selecting patients for immunotherapy treatment.

Additional studies are needed to understand if the evolution of non-BRCA1-driven HRD tumours is similar to that observed in our *Brca1^mut^* model. Studying the TIL and TME evolution in a preclinical setting where PARP inhibitors are added as a standard-of-care maintenance therapy would also be of value to mirror best the post-PARP era of HRD OC treatment.

## Data availability

Single-cell sequencing data will be made publicly available in the Gene Expression Omnibus (GEO) at the time of publication.

## Acknowledgments

We are grateful to the patients and their families for their dedicated collaboration.

We thank Jean-Paul Rivals and all the team from CHUV Biobank - Center of Experimental Therapeutics (CTE) for their assistance.

We thank the Lausanne Genomic Technologies Facility for RNA-seq analysis.

This Research Project was supported by ESMO Translational Fellowship to EG. Any views, opinions, findings, conclusions or recommendations expressed in this material are those solely of the authors and not necessarily reflect those of ESMO.

JAM-J received the support of a fellowship from “laCaixa” Foundation (ID 100010434) which code is “LCF/BQ/DR21/11880015” and a travel fellowship from the EACR.

The research was supported by the National Institute for Health Research (NIHR) Biomedical Research Centre based at Imperial College Healthcare NHS Trust and Imperial College London. The views expressed are those of the author(s) and not necessarily those of the NHS, the NIHR or the Department of Health. We thank Nona Rama, Naina Patel from the Experimental Cancer Medicine Centre (Hammersmith Campus, Imperial College) and Kay Dawson from ICHTB for tissue collection support. PC, CF acknowledge funding from the Ovarian Fund, Imperial Health Charity.

This work was supported by the Ludwig Institute for Cancer Research, the Department of Defense (DOD) Early Career Investigator (ECI) W81XWH2210703 Award OC210038 to D.D.L.

## Author contributions

E.G. and D.D.L conceptualized the study. E.G, A.J.G, N.F., M.D., F.D.C, B.S.C and H.J.F. performed the experiments. F.B., A.M., N.R., C.C., D.B, J.Do. performed the bioinformatic analysis. E.G, F.B., A.M., A.J.G, analysed and interpreted the data. J.D. provided Pathology evaluation of the samples. P.C., E.S., S.A.M., J.T., G.C., C.F. collected clinical samples. G.S., K.F., S.R., S.T. provided assistance for mIF imaging analysis. A.J.M, M.M., JA.JM., M.P., L.K., G.C., J.R.C-G. provided scientific input. E.G. and D.D.L wrote the manuscript. All authors read and approved the final version of the manuscript.

## Competing interests

C.F. received honoraria from Ethicon, GSK, Astra Zeneca/MSD, Tesaro, Clovis, Sequana and Roche, outside of the submitted work. M.M. is a current employee of the CDR-Life company. In the last three years G.C. has received grants, research support or has been coinvestigator in clinical trials by Bristol-Myers Squibb, Tigen Pharma, Iovance, F. Hoffmann-La Roche AG, Boehringer Ingelheim. The Lausanne University Hospital (CHUV) has received honoraria for advisory services G.C. has provided to Genentech, AstraZeneca AG, EVIR. Patents related to the NeoTIL technology from the Coukos laboratory have been licensed by the Ludwig Institute, on behalf also of the University of Lausanne and the CHUV, to Tigen Pharma. G.C. has previously received royalties from the University of Pennsylvania for CAR-T cell therapy licensed to Novartis and Immunity Therapeutics. JRCG has stock options with Anixa Biosciences and Alloy Therapeutics; receives licensing fees from Anixa Biosciences and consulting fees from Alloy Therapeutics; and is co-sounder of Cellepus Therapeutics. DDL has received a grant from Hoffmann-La Roche AG. All other authors declared no competing interests.

