## Supplemental Material for "Myeloid cell networks determine reinstatement of original immune environments in recurrent ovarian cancer"

- **Methods**
- **Supplemental Figures S1 to S6**
- **Supplemental Tables (zip file) Table S1 to Table S5**
- **Supplemental References**

#### Methods

##### Human ovarian cancer cohorts and samples

##### Ethics approval

This study received central approval by the University Hospital of Lausanne UNIL-CHUV and Ludwig Cancer Research Lausanne Branch institutional review board. All procedures were performed according to the Declaration of Helsinki guidelines. All cohorts were transferred to Lausanne under Standard Material Transfer Agreements for De-identified Human Tissues and Specimens Between Non-profit Organizations.

##### IMCOL primary-recurrent cohort

Forty-six patient-matched formalin-fixed paraffin-embedded (FFPE) ovarian cancer (OC) tumour samples from 23 patients were collected at the Imperial College of London at primary surgery and first recurrence. The cohort was pre-selected to be a fully platinum-sensitive cohort as reflected by the long first platinum-free interval (**Figure S1D and Table S1**). The project was performed under the Hammersmith and Queen Charlotte's and Chelsea Research Ethics Committee approval and human samples for this research project were collated by the Imperial College Healthcare Tissue Bank (ICHTB). ICHTB is approved by Wales REC3 to release human material for research (22/WA/2836) and samples were issued under full patient consent. ICHTB is supported by the National Institute for Health Research (NIHR) Biomedical Research Centre based at Imperial College Healthcare NHS Trust and Imperial College London. Samples were used for multiplexed immunofluorescence (mIF) analysis and somatic targeted DNA sequencing by ArcherDx panel.

##### UPENN primary-recurrent cohort

A total of 124 FFPE OC tumour samples (n=74 primary and n=50 recurrent samples respectively, from 46 patients) were collected at primary surgery and first recurrence under a protocol approved by the University of Pennsylvania Institutional Review Board and provided by the Tumor Tissue and Biospecimen Bank (TTAB), Department of Pathology, at the University of Pennsylvania, Philadelphia, USA. Detailed clinical data and samples information are reported in **Table S1.** Sample were used for mIF analysis and somatic targeted DNA sequencing by ArcherDX panel.

##### HiTide-UPENN primary cohort

Fifty-one FFPE samples, n=71 snap-frozen samples, and n=19 freshly cryopreserved tumour samples were collected from 51 patients at the Ovarian Cancer Center, Department of Obstetrics & Gynecology, University of Pennsylvania, Philadelphia, USA. Informed consent was obtained from all subjects included in this study under an approved protocol from the Institutional Review Board (UPCC 17909, IRB 702679) under the care of Dr DJ Powell and Dr J Tanyi. Samples were collected from unselected consecutive patients ("all comers") undergoing surgery for primary stage III or IV high-grade serous ovarian cancer, as well as from the fallopian tube or primary peritoneal origin. Detailed clinical data are reported in **Table S2**. Samples were used for mIF analysis, bulk RNAsequencing, somatic DNA sequencing by the BROCA panel (performed by Dr E. Swisher) and MultiNicheNet analysis on single-cell RNAsequencing.

**FFPE slides preparation and mIF staining**

Slides were prepared at the Immune Landscape Laboratory (ILL) at the Center for Experimental Therapeutics (CTE) of the Department of Oncology at CHUV (Lausanne, Switzerland) from the FFPE blocks provided.

The first slide was used for H&E staining and review by a dedicated Pathologist (JD) to define the quality of tumor and stromal areas and exclude adjacent healthy tissue. A second slide of 3.5 um was used for mIF. Slides to be stained were thawed and heated at 60°C for 1 hour. mIF panels were run using the multiplex Ventana Discovery ULTRA Staining module autostainer (Roche). Slides were placed on the staining module for the deparaffinization step, consisting of 3 cycles of 8 minutes at 69°C (Discovery Wash,Ventana Roche), followed by epitope retrieval for 64 minutes at 95°C or 98°C (according to the panel protocol performed) in high pH buffer Cell Conditioning 1 (CC1, Ventana Roche) and endogenous peroxidase quenching (Discovery Inhibitor, Ventana Roche). The automated immunofluorescence (IF) staining procedure consists of multiple consecutive rounds (6 for a 7-plex) of staining. Each round includes non-specific sites blocking (Ventana, Discovery Inhibitor and Discovery Goat Ig Block), incubation with unlabeled primary antibody, followed by incubation with horseradish peroxidase (HRP)-conjugated secondary antibodies (Discovery OmniMap anti-Rabbit (Rb) and anti-Mouse (Ms), Ventana), and OpalTM (Akoya) reactive fluorophore (Opal 480, 520, 570, 620, 690, 780) detection that covalently labels the primary epitope. Then an antibody (both primary and secondary) heat denaturation step was performed prior to the next round of antibody staining. Finally, Spectral DAPI (Akoya) was used for nuclear staining.

The following mIF panels have been applied:

1) CD8 (rabbit, @CellMarque clone 108R-16-RUO), CD11c (mouse, @CellMarque clone 5D11), PD1 (mouse @BioCare clone ACI3137C), PD-L1 (rabbit, @CellSignaling, clone E1L3N 13684S), CD68 (anti-mouse, @DAKO clone PG-M1), pan-Cytokeratin (anti-mouse @Dako clone M3515), DAPI

2) CD3 (rabbit, @DAKO clone A0452), CD8 (rabbit, @CellMarque clone 108R-16-RUO), PD1 (mouse, @BioCare clone ACI3137C), GzB (mouse, @Monosan clone MON7029-1), CD103 (rabbit, @CellMarque clone EP206), pan-Cytokeratin (anti-mouse @Dako clone M3515), DAPI.

mIF images were acquired on the Vectra® Polaris automated quantitative pathology imaging system (Akoya Biosciences). This multispectral imaging system uses the MOTiF technology, allowing the unmixing of spectrally overlapping fluorophores and tissue autofluorescence of whole slide scans. For the optimal IF signal unmixing (individual spectral peaks) and the subsequent multiplex analysis, a spectral library containing the individual emitting spectral peaks of all fluorophores was created. For this, single antibody-coupled with fluorophore staining under the optimized conditions without DAPI, and a DAPI single staining, were performed. In addition, auto-fluorescence controls were performed by staining tumor tissue slides omitting both the fluorophore and DAPI.

**mIF data analysis**

**Cell-densities and Immune-classification algorithm**

Using the Phenochart™ whole-slide viewer, regions of interest (ROIs, 931um x 698 um, range 5-110 ROIs per section) representative of the entire FFPE tissue sample were acquired. InForm 2.5.1 (Akoya Biosciences) software was used for training and phenotyping analysis. The images were first segmented into specific tissue categories of tumour, stroma and no tissue, based on pancytokeratin (panCK+) and DAPI staining using the Inform Tissue Finder™ algorithm. Individual cells were then segmented using the counterstained-based adaptive cell segmentation algorithm. Quantification of the immune cells was then performed using the Inform active learning phenotyping algorithm by assigning the different cell phenotypes across several images chosen for the project. IF-stained cohorts were then batch processed and data were exported via an in-house developed R-script algorithm (Post-InForm) to retrieve single-cell x,y coordinates and staining positivity.

To calculate cell densities, we counted the number of a specific cell phenotype in both tumour and stromal compartment across the whole FFPE tissue section. Counts of each specific cell type (tissue-specific) were then divided by the area of the tissue ($mm^2$) to obtain a density (number of cells/$mm^2$).

We then developed a two-step algorithm to define a $CD8^+$ T cells-based immune classifier considering not the average cell density across the whole tissue but the heterogeneous $CD8^+$ T cells distribution within tumour/stroma compartments of each ROI. First, we computed the fraction of inflamed subregions (ROIs with >21 $CD8^+$cells/$mm^2$) in tumour and named the sample purely-inflamed if the percentage of inflamed ROIs was >70% and as mixed-inflamed when this percentage was between 50% and 70%. Second, in case less than 50% of ROIs were inflamed in the tumour, we considered the sample as excluded if >10% of ROIs showed >21 $CD8^+$ cells/$mm^2$ in the stroma and as desert if this percentage was <10%.

**Neighborhood mutual cell interaction analysis**

Starting from previous published works[1,2] we computed a new "mutual interactions" methodology which takes into consideration "bi-directional" cells interaction from two different cell types. Normalization by the proportions of both cell types of interest on a surface (i.e. tumour or stroma regions) was applied to avoid cell abundance bias.

In our specific case, the total area, can be split into tumoural regions (defined by the presence of the $panCK^+$ protein marker) and stromal regions (defined by the absence of $panCK^+$) where we can measure the neighboring of a point (i.e. cell) (defined as starting point) of type "A" (i.e. cell type A) of coordinates (xa, ya), with type "B" points (i.e. cells from cell type B; defined as ending points) in the surrounding area within a predefined distance (i.e. 20um). The distance between point "A" (i.e. cell type A) and a point "B" (i.e. cell type B), with coordinates (x, y), is then defined as the euclidean distance D:

$$D = \sqrt{(x_a - x_b)^2 - (y_a - y_b)^2}$$

A point "A", and a point "B" are then considered neighbors, if the distance "D", between "A" and "B" is less than a given threshold (ε). Such definition can be visualized as a circular surrounding of the point A with a radius equal to the threshold (ε, that is 20 um in our case).

In particular, given two sets of points in a two-dimensional space we could implement a measure that estimate their vicinity. This measure is called mutual interaction [3].

Given the set of points (i.e. cells) A={A1, …., AN}, and the set of points (i.e. cells) B={B1, …, BK}, for each point of the set "A" we measure if there is, at least, an element of the set "B" at a distance D< ε. For a given point, if the condition is true, such a point has at least one neighbor of type "B". We then sum all the points of the set A that have at least one neighbor of type "B". We repeat such procedure, but now starting from the points of the set "B" looking for neighbors in the set "A".

The mutual interaction is them computed as:

(number of A with a neighbor B) + (number of B with a neighbor A) /( Number of A + Number of B).

**Mathematical Representation of Mutual Interaction in Sets A and B**

In our methodology, we introduced a mathematical expression to quantify the mutual interaction between sets A and B. This expression captures the condition that an element in one set has at least one neighbor in the other set. We denoted this condition using the underscore symbol (_).

Let: N(A_B) be the number of elements in set A that have at least one neighbor in set B,

N(B_A) be the number of elements in set B that have at least one neighbor in set A,

N(A) be the total number of elements in set A,

N(B) be the total number of elements in set B.

The mutual interaction measure is expressed as:

$$M = \frac{N(A\_B) + N(B\_A)}{N(A) + N(B)}$$

**Tissue Partitioning for Mutual Interaction Computation**

In the context of tissue partitioning, the mutual interaction value is computed separately for cells in the tumour and stroma regions. This process involves considering points in the stroma (or tumour) from set "A" and computing the neighboring metric with all points in set "B" (not restricted to stroma only). Similarly, the measure is repeated for points in the stroma (or tumour) from set "B," computing the neighboring metric using all points in set "A."

To ensure comparability, the computed values are normalized by the sum of points in sets "A" and "B" in stroma (or tumour). This normalization accounts for the varying cell densities in different tissue compartments, providing a more accurate assessment of mutual interactions.

This approach allows for a nuanced understanding of cellular dynamics within specific tissue environments, considering the unique interactions occurring in both tumour and stroma regions.

**Cellular Triplets Spatial Interactions**

Building upon the established function for pairwise interactions, our methodology extends seamlessly to analyze cellular triplets. Consider three sets of points: "A," "B," and "C," each representing distinct cell types.

For triplets, the mutual interaction is computed as follows:

$$M = \frac{N(A\_B\_C) + N(B\_A\_C) + N(C\_A\_B)}{N(A) + N(B) + N(C)}$$

**Survival curves**

Survival curves for overall survival (OS) in our clinical cohorts were constructed using the Kaplan–Meier estimator and statistical significance was determined using the log-rank test.

**Targeted DNA analysis**

**UPENN and IMCOL primary-recurrent cohorts:**

**DNA extraction and libraries preparation**

Three sections of 8um-thick were freshly cut from each FFPE blocks. Tissues were deparaffinized with 500ul Deparaffinization solution (Qiagen) and total DNA was extracted with QiAmp DNA FFPE tissue kit (Qiagen). Final DNA was eluted in 25ul of ATE buffer. DNA concentration was measured with Qubit DNA fluorometer method (inVitrogen). A minimum of 80 ng of sample input (100ng preferred) was used to construct the targeted libraries with Archer VariantPlex somatic protocol for Illumina, following manufacturer's instructions. Reagents were supplied by ArcherDX, including our custom panel of 49 gene-specific primers that target regions of interest and Archer MBC adapters to tag each unique molecule with a barcode. Final libraries were quantified by Qubit DNA method and quality was checked on Fragment Analyzer (HS-NGS fragment kit, Agilent).

**Sequencing and analysis**

Libraries were pooled at equimolar concentrations and loaded into the Miseq or Novaseq Illumina system for sequencing, following ArcherDX recommendations. Fastq files generated were analyzed using Archer analysis software. The reads are aligned with BWA-MEM and PCR duplicates are removed. Single nucleotide variants and indels are identified using HaplotypeCaller for both tumour and three unmatched normal tissue samples. The mutations in tumour samples are cleaned by removing mutations that are also present in normal samples. Total read depth and variant allele frequency filters are applied to remove potential artifacts. Ambiguous mutations are cross checked using Archer mutation calling pipeline and only kept if they were reported in both pipelines. Mutations are annotated with ClinVar to determine pathogenic variants. For identifying DSB repair pathway mutations only the truncating mutations, pathogenic mutations or damaging mutations according to both SIFT and PolyPhen are considered.

**HiTide-UPENN primary cohort:**

Mutations in the TP53, BRCA1, BRCA2 and RAD51C genes and methylation of BRCA1 were identified as previously described[4-6] through BROCA targeted panel.

**Bulk RNA sequencing library preparation and processing**

Total RNA was extracted from snap frozen tissues. Tumour tissues were disrupted on ice in RLT buffer supplemented with ~40mM dithiothreitol (DTT, Sigma Aldrich), using a pestle (70 mm, 1.5/2.0 mL, Schuett-Biotec). Lysates were further homogenized using a syringe and needle. After centrifugation at full speed for 3 min in a benchtop centrifuge (Eppendorf) at 4 °C, supernatant was used for RNA extraction according to manufacturer's protocol, including on column DNase digestion, using the RNeasy Mini Kit (Qiagen). RNA quality was assessed with a Fragment Analyzer (Agilent) and Nanodrop One spectrophotometer (Thermo Scientific). Quantification was performed with the Qubit RNA broad-range (BR) assay kit (Invitrogen). RNA sequencing libraries were prepared using the Illumina TruSeq Stranded RNA reagents according to the protocol supplied by the manufacturer and sequenced using HiSeq 4000/Novaseq. Illumina paired-end sequencing reads were aligned to the human reference

GRCh37/hg19 genome using STAR aligner (version v2.7.3a; https://github.com/alexdobin/STAR) and the 2-pass method as briefly follows: the reads were aligned in a first round using the --runMode alignReads parameter, then a sample-specific splice-junction index was created using the --runMode genomeGenerate parameter. Finally, the reads were aligned using this newly created index as a reference. To transform raw counts into TPM values, raw counts were summarized at the gene level using htseq-count (version 0.9.1). Read counts where then normalized into reads per million (TPM). The comprehensive gene annotation version 32 was downloaded from the GENCODE website (https://www.gencodegenes.org/human/release_32lift37.html) and chromosome position, transcript structure and transcript and protein sequences were selected to annotate genes.

**Gene expression analyses**

Genes with zero expression and with low gene count variance were filtered out from the analysis and transcript per million, TPM, values were used for the downstream analysis. Additionally, genes sharing the same enzymatic function were collapsed using the geometric mean. Analysis was performed in R language for statistical computing. Gene set variation analysis enrichment scores were calculated using GSVA R package and subsequently clustered using euclidean distance and ward.D2 method in with the pheatmap R package. The p-values in the boxplots were calculated using Wilcoxon test and were adjusted with the Bonferroni correction.

**Gene signatures**

The detailed description of the gene sets is provided in **Table S2**. More than half of the signatures were derived from the MSigDB database Hallmark collection (full) and C2 collection (selected signatures). About 10% of the signatures were compiled from the important signatures previously identified in our Lab (complete references are indicated in the table).

**Single-cell RNA sequencing (HiTide-UPENN cohort)**

Vials of dissociated cells cryopreserved in 90% Human Serum + 10% dimethyl sulfoxide (DMSO) were used as starting material. The day of the assay, cells were thaw in RPMI + 10%FBS and manually counted on a hemocytometer using Trypan blue (Gibco) exclusion. Two to four million of cells were then FcR blocked (Miltenyi Biotec) for 15min at RT. After incubation and washing, cells were stained with CD45-APC (#304012, BioLegend) for 20min at 4°C and resuspended in PBS + 0.04% BSA (Sigma-Aldrich) + 0.1% RNasin (Promega) for sort. DAPI staining (Invitrogen) was performed for 1min at RT.

**FACS sorting, encapsulation and library construction**

For each sample, 40’000 CD45+ cells were sorted on a MoFlo Astrios (Beckman Coulter) and collected in a 0.2mL PCR tube containing 10µl in PBS + 0.04% BSA + 0.1% RNasin. After sorting, cells were manually counted with a hemocytometer and viability was assessed using Trypan blue exclusion. Single-cell RNA libraries were generated using the Chromium Next GEM. Single Cell 5’ Library and Gel beads kit v2 (10X Genomics). Cells were resuspended at a density of 600-1200 cells/µl with a viability >90% for single-cell analysis. When possible, 15’000 cells of each sample were loaded into the Chromium machine, encapsulated and barcoded following the manual CG000207 from 10X Genomics (5’ chemistry). All library construction steps were performed according to the manufacturer's protocol. Complementary DNA and library quality were examined on a Fragment Analyzer (Agilent) and quantification was performed with the Qubit HS dsDNA assay kit (InVitrogen).

**Single-cell RNA aligning and downstream analysis**

Alignment, barcode and UMI counting were performed using GRCh38-2020-A reference genome and cellranger-6.1.1 from 10x Genomics. Multiple gene expression libraries were combined via *cellranger aggr* and filtered feature-barcode matrix containing gene expression data from 3’ libraries was further analyzed with the Seurat R package. The annotation of the cell states and lineage assignments were performed as described in detail below for the mouse single cell sequencing analysis with some minor variations. For example, *infercnv* algorithm in the human annotation case was applied to all cells and cells with high number of copy number changes were assigned to malignant lineage and removed from downstream analysis.

**Mice**

**Cell cultures**

ID8 *Trp53*$^{-/-}$*Brca1*$^{wt}$ and *Trp53*$^{-/-}$*Brca1*$^{mut}$ mouse OC cell lines, obtained from the laboratory of Prof. Iain A. McNeish (Institute of Cancer Sciences, University of Glasgow, Scotland)[7,8] were transduced to express luciferase [1] and cultured in DMEM supplemented with 4% FBS, 100 µg/mL penicillin, 100 µg/mL streptomycin, and ITS 35 (5µg/mL insulin, 5µg/mL transferrin, and 5ng/mL sodium selenite). Cell lines were negative for Mycoplasma contamination.

**Model**

C57BL/6NHsd female mice were obtained from Envigo and were maintained in pathogen-free conditions. Age-matched mice 7 weeks were used for all experiments. Animal experimentation procedures were performed according to the protocols approved by the Veterinary Authorities of the Canton Vaud (VD3480d and VD3480x1b), according to Swiss law.

We injected 5 × 10^6 ID8 derivative cancer cells expressing luciferase (ID8Luc) i.p. in C57BL/6NHsd female mice. To mimic OC standard of care we treated mice with i.p. Carboplatin (Accord, ref: 7504554, 20mg/kg) – Taxol (Labatec, ref: 4670594, 3mg/kg) once weekly for 6 weeks, reach tumour control (luciferase signal, SD) and then wait for tumour recurrence.

Mouse health and welfare were monitored regularly. For both control groups and experiments evaluating survival post-therapy, we used body and health performance score sheets (taking into consideration ascites accumulation) and mice were sacrificed once reaching the equivalent of humane endpoints.

**Abdominal Bioluminescence imaging**

Tumor growth was monitored by Bioluminescent imaging (BLI). BLI was performed using the Xenogen IVIS® Lumina II imaging system and the photons emitted by the Luciferase-expressing cells within the animal body were quantified using Living Image software. Briefly, mice bearing ID8Luc cancer cells were injected i.p. with D-luciferin (150mg/kg stock, 100 µL of D-luciferin per 10 g of mouse body weight) resuspended in PBS and imaged under isoflurane anesthesia after 10 min. A 49 pseudocolor image representing light intensity (blue, least intense; red, most intense) was generated using Living Image. BLI findings were confirmed at necropsy.

**Tumour processing and flow cytometry**

At the time of sacrifice, i.p. tumours were dissected. Tumours were digested in 200 µg/ml Liberase TL and 5 units/ml DNase I in DMEM for 30min at 37°C, with rotation. For ex vivo staining, 1-2x$10^6$ cells were stained with LIVE/DEAD Fixable Aqua Dead Cell Stain (1:500). Fc receptors were blocked for 10 min at 4°C with 5 µg/ml Mouse BD FC Block. Cells were fluorescently labeled with antibodies for 30 min at 4°C, washed and resuspended in fixation buffer (2% formaldehyde in PBS) or intracellularly stained according to the manufacturer's protocol (eBiosciences). Flow cytometric analysis was performed on LSR II flow cytometer and analyzed using FlowJo software.

**Single-cell RNA sequencing**

Cells were counted on the ADAM automated cell counter and viability was estimated with the AccuStain solution kit (NanoEntek). Cells were surface-stained with CD45-BV785 + CD8-BU650 for 20min at 4°C and resuspend in 1ml of PBS+0.04% BSA (Sigma-Aldrich) after washing.

**Cell multiplexing**

After staining, samples were labeled and multiplexed by group, allowing to be pooled in a single GEM for encapsulation. Cell labeling was performed according to the Cell multiplexing oligo labeling protocol from 10x Genomics (CG00391). 500'000 cells per sample (if possible, otherwise minimum 200'000 cells) were labeled with a cell multiplexing oligo for 5min at room temperature. After two washes with PBS+ 1% BSA, cells were resuspended in PBS + 0.04% BSA + 0.1% RNasin and multiplexed by equimolar pools. Before sorting, 10min of viability staining with Reddot1 (Biotium) and 3min of DAPI staining were performed.

**FACS sorting**

50'000 total live cells were sorted for each pool on a MoFlo Astrios (Beckman Coulter) and collected in 0.2mL PCR tubes containing 10ul in PBS + 0.04% BSA + 0.1% RNasin. After sorting, cells were manually counted with hemacytometer and viability was assessed using Trypan blue exclusion.

**Encapsulation and library construction**

Single-cell RNA libraries were generated using the Chromium Next GEM Single Cell 3' Library and Gel beads kit v3.1 according to the manufacturer's instructions. For each sample, 15'000 to 30'000 cells were loaded into the Chromium machine, encapsulated and barcoded following the manual (CG000388), aiming a recovery of 10'000 to 20'000 cells according to manufacturer conditions. After encapsulation and reverse transcription, 11 PCR cycles were used to amplify cDNA. All libraries construction steps were performed according to the manufacturer's protocol. For each sample, 3GEX library was generated. If sample was part of a pool, a Cell Multiplexing Library (CML) was also constructed. Complementary DNA and library quality were examined on a Fragment Analyzer (Agilent) and quantification was performed with the Qubit HS dsDNA assay kit (InVitrogen).

**Sequencing**

Barcoded 3'GEX libraries and CML were pooled and sequenced on an on Illumina HiSeq 4000 or NovaSeq6000 system, following 10X Genomics recommendations. GEX libraries were sequenced to a median depth of 20,000 unique reads per cell and Cell Multiplexing Oligos (CMO) libraries were sequenced to a median depth of 5'000 unique reads per cell.

**Alignment, annotation and downstream analysis**

Alignment, barcode and UMI counting were performed using mm10-2020-A reference genome and cellranger-6.1.1 multi from 10x Genomics. Multiple gene expression libraries were combined via *cellranger aggr* and filtered feature-barcode matrix containing gene expression data was further analyzed with the Seurat R package. The total number of cells detected was 50949, with number of cells successfully assigned to an individual mouse CMO library ranging between 15% and 85%. To rescue the cells that were not assigned to any CMO library (i.e. to any individual sample), the alignment procedure from above was repeated in *cellranger* without providing the CMO library information. Next, all the cells whose barcodes were not mapped to any of the CMO libraries, were pooled together for each corresponding mouse group as a pseudo-mouse and added to the resulting matrix for downstream analysis.

For annotation purposes, several iterations were performed. In the first iteration, cells were clustered at a high resolution leading to a big number of cell clusters with shared properties. The clusters were obtained using the standardized Seurat procedure: data counts were log normalized using the *NormalizeData* function, then variable features were found using *vst* method and 600 fetures. Next, a linear transformation using the *ScaleData* function was applied, and linear dimensional reductions were calculated using *RunPCA* (for the principal component analysis) and *RunTSNE* (for the t-Distributed Stochastic Neighbor Embedding) functions with first ten principal components used as input features and perplexity of 30. Finally, shared nearest neighbors, SNN, was calculated using *FindNeighbors* (with 10 PCs and k=30) followed by *FindClusters* functions (with resolution = 20).

Based on the known exclusive markers, the cells were automatically classified as immune (*Cd3g*, *Cd3d*, *Cd3e*, *Cd2*, *Cd8a*, *Cd8b1*, *Foxp3*, *Il2ra*, *Trbc1*, *Cd19*, *Ms4a1*, *Cd79a*, *Cd79b*, *Ncr1*, *Klrb1c*, *Klrd1*, *Klrk1*, *Aif1*, *Ms4a7*, *Cd14*, *Fcgr4*, *Itgam*, *Itgax*, *Mrc1*, *Cd163*, *Fcer1a*, *Clec10a*, *Mzb1*, *Derl3*), non-immune (*Epcam*, *Msln*, *Egfr*, *Fap*, *Pdpn*, *Dcn*, *Thy1*) or endothelial cells (*Pecam1*, *Fas*) if at least 80% of the cluster's cells expressed the markers. The cells within these 3 initial categories were further automatically classified as T cells (*Cd3g*, *Cd3d*, *Cd3e*,

*Cd2*, *Cd8a*, *Cd8b1*, *Foxp3*, *Il2ra*, *Trbc1*), B cells (*Cd19*, *Ms4a1*, *Cd79a*, *Cd79b*), NK cells (*Ncr1*, *Klrb1c*, *Klrd1*, *Klrk1*), Myeloid cells (*Aif1*, *Ms4a7*, *Cd14*, *Fcgr4*, *Itgam*, *Itgax*, *Mrc1*, *Cd163*, *Fcer1a*, *Clec10a*, *Mzb1*, *Derl3*, *Cd48*), endothelial cells (*Pecam1*, *Fas*), malignant (*Epcam*, *Msln*, *Egfr*, *Brca1*, *Brca2*, *Trp53*), fibroblasts (*Fap*, *Pdpn*, *Dcn*, *Thy1*, *Ankrd1*, *Mcam*, *Cd70*, *Pdgfra*, Pdgfrb, *Itga5*, *Mme*) if at least 50% of the cells expressed the markers. At the end of the initial classification, the expression of the above markers along with the additional list of markers was visualized using the *doHeatmap* function in each of the assigned classes to verify the validity of classifier.

After the first iteration, a filtering step was applied individually on each of the identified lineage depending on the total distribution of that lineage population: number of genes from 250 to 3000-6000; number of reads 500 to 15000-40000; below 15% mitochondrial content and within 1-7.5 to 40% ribosomal content. This reduced the total number of cells by 15%.

At the second iteration, for the filtered cells from each individual library, variable genes from the log-normalized counts were found using vst method and then the libraries were integrated using the anchoring technique described in "Stuart and Butler et al"[9]. As during the first iteration, integrated data was scaled and then passed to PCA, t-SNE, and SNN analyses for identifying clusters (with resolution = 0.3). Next, gene expression centroids (average gene expression profiles per cell type) method was applied using matrices from Zilionis et al. [10] for main and sub-populations to predict the cell type of each given cell. Following the centroids methods prediction, the cell annotation was refined per each individual cluster using its initial assignment, its predicted state, and expression of a particular known markers (like *Cd8* for Cd8$^+$ T cells and *Foxp3* for Tregs). This refinement completed the second iteration of the cell annotations.

At the last iteration, based on the annotations obtained from second iteration, cells were divided and re-clustered in five main groups or lineages: T cells, B cells, Myeloid, Malignant and doublets. For each of the main lineages of cells, the CMO associated genes were filtered out from the analysis and the normalization to clustering (resolution = 0.3) steps were performed as described above. Then, differentially expressed genes (using the *FindAllMarkers* function) were identified for each cluster and signature scores were calculated for each cell using the *AUCell* R package and "in-house" signatures defined in **Table S2**, Additionally, centroids method was applied to predict the states from Barras et al.[3] for each cell. Once all the above metrics were calculated, the cells per each cluster were manually refined considering all the newly obtained metrics and the initial annotation. When necessary, clusters were re-assigned to the different main lineage to reflect the observed DE genes in these cluster and signatures expressed in corresponding cells. Also, due to the gene expression dropout issue, for the cells that clearly expressed T cell markers, but were double negative for *Cd8* and *Cd4* expression, the assignment to *Cd8* versus *Cd4* group was based on predicted value from the centroid algorithm and based on which group of cells they clustered with. In addition, when markers from multiple lineages were expressed on the same cells (e.g. $Cd8^{+}Cd79a^{+}$ cells), these cells were categorized as doublets and omitted from the downstream analysis.

Finally, the copy number inference algorithm from *infercnv* R package was applied to all malignant cells to validate their malignant state. A total of 900 cells (300 cells each) randomly selected from T, B and myeloid compartments were used as a normal reference to infer the copy number changes of the tumor cells. The low number of the resulting inferred copy number variations in the normal cells and high number of those in tumor cells validated the correctness of malignant cell assignment.

The images for the downstream analysis were produced either by the built-in functions from Seurat package or by ggplot2 package.

**MultiNicheNet analysis**

MultiNicheNet (MNN) (version 1.0.3) was utilized to explore the differences in ligand-receptor interactions. Throughout our analysis, we used the default parameters to look at the top 250 targets with minimum log-fold change of 0.5 and a fraction cut-off of 0.05.

For the ligand-receptor analysis in the human cohort (**Figure 3J**) we analyzed N=76'586 cells (31'293 of which were either $CD8^{+}$ T cell, $CD4^{+}$ T cell, macrophages or dendritic cells) from N= 19 samples (N=4 inflamed, N= 11 excluded, N= 4 desert).

We separated the data into immune phenotypes and restricted it to four major cell lineages: $CD8^{+}$ T-cells, $CD4^{+}$ T-cells, macrophages and dendritic cells. Due to the low number of samples by immune phenotype and by following authors recommendations[11], we decided to utilize empirical p-values (empirical_pval = TRUE) and use the normal p-values to retain differentially expressed genes.

For the ligand-receptor analysis in the mouse dataset, we initially encompassed the major cell types involved (**Table S5):** B cells, CD8 and CD4 T cells, DC cells, macrophages, malignant cells and stromal cells from N=12 recurrent samples (6 *Brca1*$^{mut}$ and 6 *Brca1*$^{wt}$) for a total of

23’449 cells (12’595 in the *Brca1*$^{mut}$ and 10’854 in the *Brca1*$^{wt}$). Subsequently, we conducted a more focused investigation on the above samples targeting the most promising interactions from malignant cells, macrophages, and DCs cells, considering finer annotation (N=11’978 total cells, 5’379 in the *Brca1*$^{mut}$ and 6’599 in the *Brca1*$^{wt}$ respectively). Due to a higher abundance of samples per category, we maintained recommended parameters such as "adjusted p.value = TRUE" "empirical_pval = FALSE". We used the get_top_n_lr_pairs function to generate two key outputs: the top 50 scaled products of ligand and receptor expression within each experimental group, enabling us to estimate their ligand activity (**Figure S6B**) and a regulatory network highlighting the top 150 ligand-target gene interactions (**Figure S6A**). For visualization, we specifically selected the 50 best-predicted interactions (**Figure 6E**).

**Multiplex chromogenic Immunohistochemistry**

The triple chromogenic immunohistochemistry assay was performed using the Ventana Discovery ULTRA automate (Roche Diagnostics, Rotkreuz, Switzerland). All steps were performed automatically with Ventana solutions except if specified otherwise. Dewaxed and rehydrated paraffin sections were pretreated with heat using the CC1 solution for 40 minutes at 95°C. Primary antibodies were applied and revealed sequentially either with a rat Immpress HRP (Ready to use, Vector laboratories Laboratories) or a rabbit UltraMap HRP followed by incubation with a chromogen (ChromoMap DAB, Discovery purple and Discovery Teal). A heat denaturation step was performed after every revelation. The primary antibodies sequence was: rat anti-CD11c, rat anti-CD8 and rabbit anti-PanCytokeratin. Sections were counterstained with Harris hematoxyline (J.T. Baker) and permanently mounted with Pertex (Sakura). For immunohistochemical quantification of $CD8^{+}$ cells and $CD11c^{+}$ cells, 10 × 10 tiled bright-field pictures of FFPE sections were taken at 100 µm magnification to cover almost whole slide surface. Cell counts were obtained using ImageJ software.

### Supplemental Figures

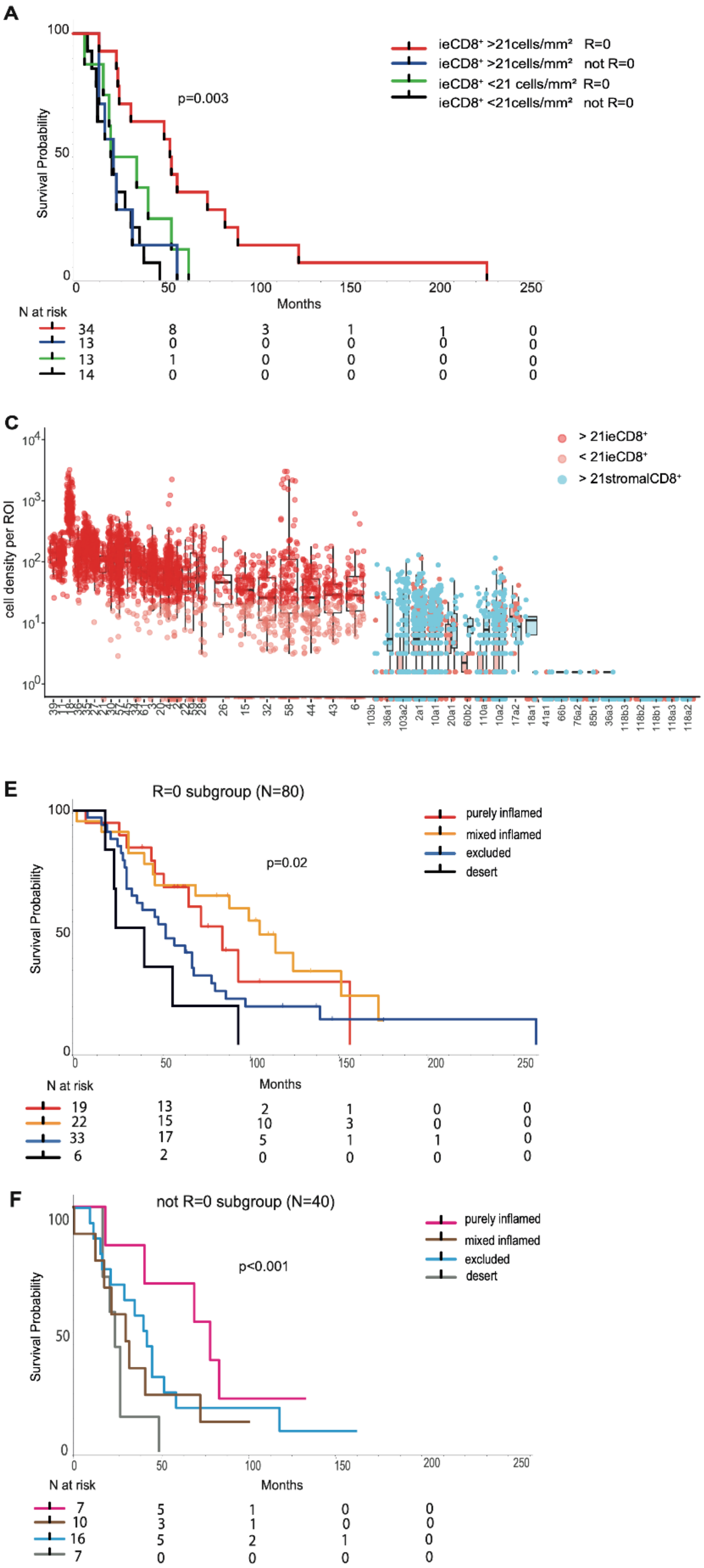


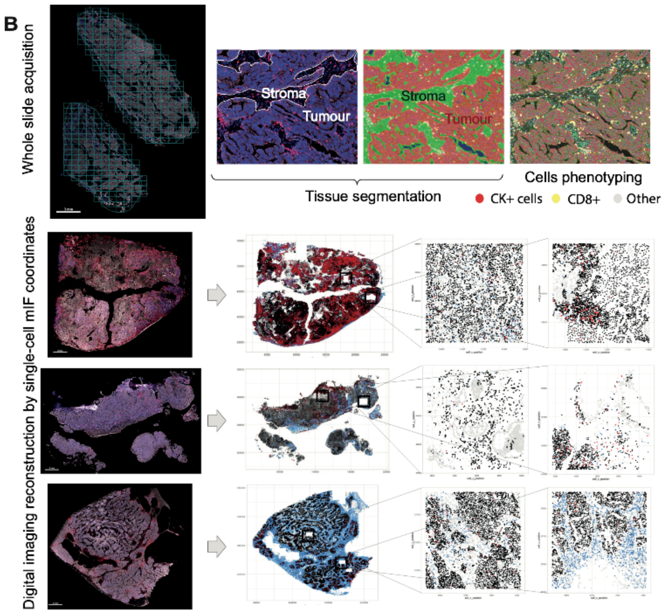


D

| Characteristic primary cohorts | IMCOL (N=23) | UPENN (N=74) | UPENN-HiTide (N=51) | |
|---|---|---|---|---|
| **Median age at diagnosis** (years, range) | 59 (44-71) | 54 (25-81) | 59 (36-78) | |
| **Histology** (N, %) | | | | |
| Serous | 20 (87%) | 48 (89%) | 45 (88%) | |
| Other | 3 (13%) | 4 (7%) | 6 (12%) | |
| unknown | - | 2 (4%) | - | |
| **FIGO stage** | | | | |
| ≤IIc | **8 (35%)*** | 4 (7%) | - | **p=0.0003** |
| IIIc | 14 (61%) | 42 (77%) | 46 (90%) | |
| IV | 1 (4%) | 6 (11%) | 5 (10%) | |
| unknown | - | 2 (4%) | - | |
| **Residual disease at 1st surgery** | | | | |
| R=0 | 20 (87%) | **38 (70%)*** | 43 (84%) | **p=0.0006** |
| Not R=0 | 3 (13%) | 14 (26%) | 7 (14%) | |
| unknown | - | 2 (4%) | 1 (2%) | |
| **Median PFS** (months, range) | 29 (13-99) | 10 (4-84) | N/A | |

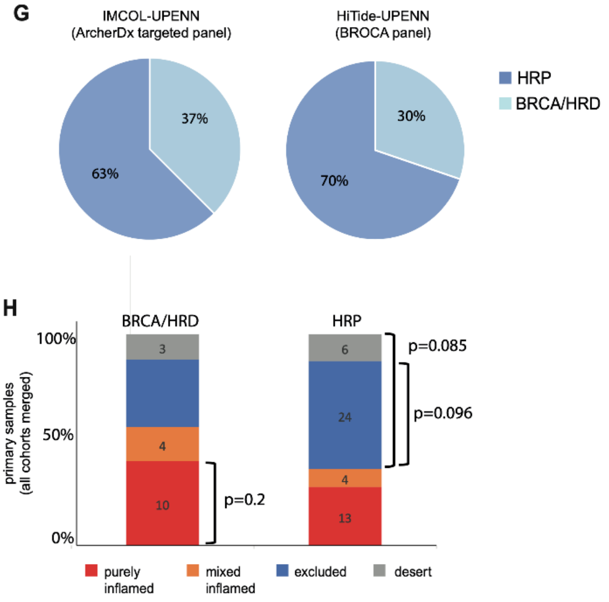

**Figure S1: Whole-slide CD8$^+$ multiplexed imaging and CD8$^+$ based immune classification of OC**.

**(A)** Kaplan-Meyer curves of overall survival (OS) in the IMCOL and UPENN cohort merged according to the new mIF cut-off of ie21CD8+/mm$^2$ (as a median on whole slide) and the absence of residual disease (R=0 or not) at primary surgery (p value assessed using the Log-rank test). **(B)** Schematic representation of the mIF analysis process: acquisition of multiple equal subregions or ROIs to cover all slide, tissue segmentation in tumour/stroma regions according to panCK$^+$ expression, cell phenotyping (InformTm and Methods), example of digital imaging reconstruction using single-cell coordinates (X,Y) and representative example of regions of interest (ROIs) extraction from the original mIF image. **(C)** Representative miF images examples of the four CD8 immune categories. **(D)** Selected clinical baseline characteristics among the three different OC cohorts (extended data reported in Supplementary Table 1 and 2). **(E-F)** Kaplan-Meyer curve of OS merging the three cohorts in the optimal cytoredution (R=0) and not-optimal (not R=0) subgroups respectively according to immune category (p value assessed using the Log-rank test). **(G)** Pie-charts showing the percentage of HRD/BRCA tumours in the IMCOL-UPENN cohorts and HiTide-UPENN. **(H)** Bar-plots showing the proportion of immune-categories in the BRCA/HRD and HRP subgroups in primary tumors (percentage in the y axis and exact numbers reported within the bar-plots); p-value assessed using the hypergeometric test.

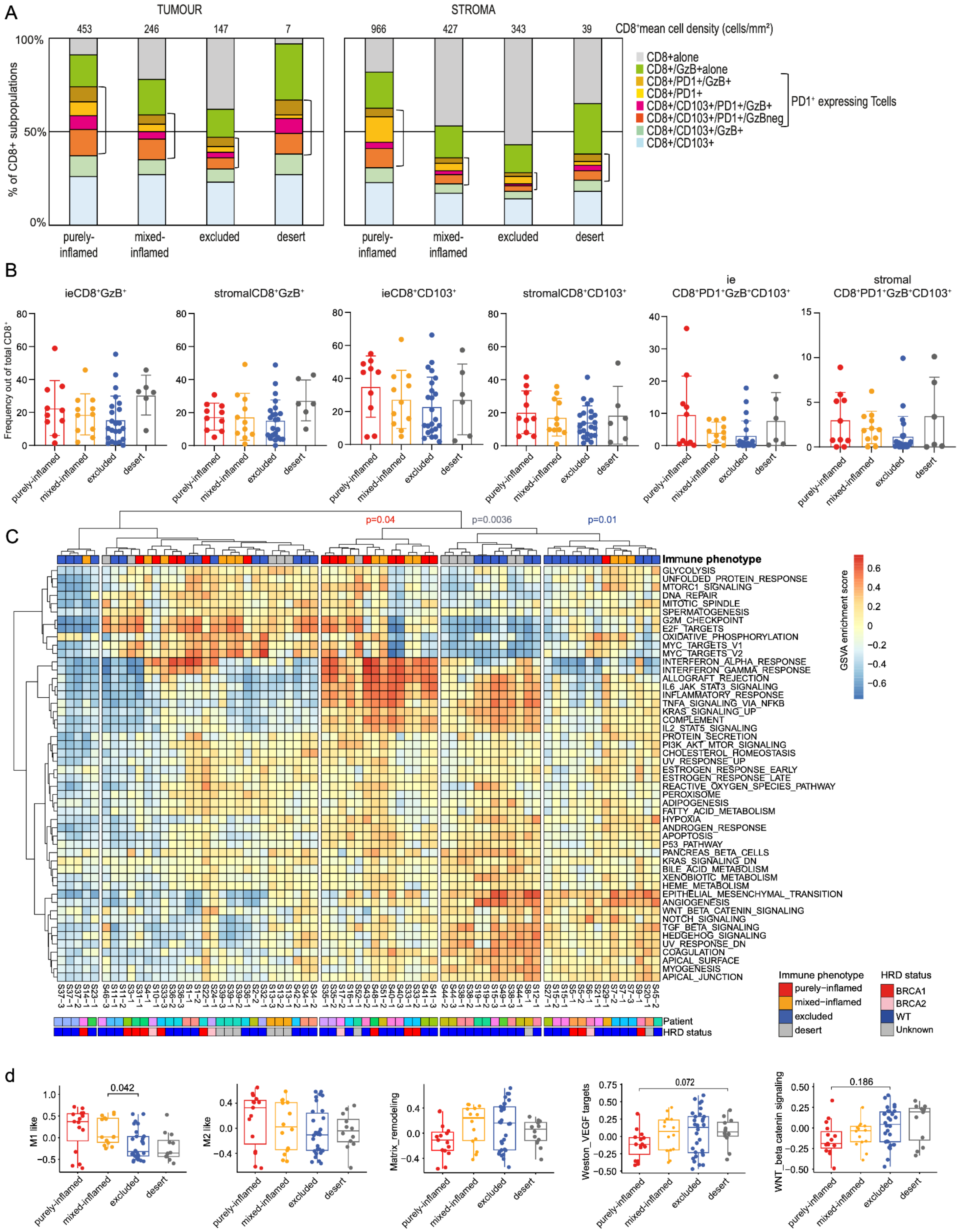

A
TUMOUR
STROMA
CD8+mean cell density (cells/mm²)
453
246
147
7
966
427
343
39
% of CD8+ subpopulations
100%
50%
0%
purely-inflamed
mixed-inflamed
excluded
desert
CD8+alone
CD8+/GzB+alone
CD8+/PD1+/GzB+
CD8+/PD1+
CD8+/CD103+/PD1+/GzB+
CD8+/CD103+/PD1+/GzBneg
CD8+/CD103+/GzB+
CD8+/CD103+
PD1+ expressing Tcells
B
ieCD8+GzB+
stromalCD8+GzB+
ieCD8+CD103+
stromalCD8+CD103+
ie CD8+PD1+GzB+CD103+
stromal CD8+PD1+GzB+CD103+
Frequency out of total CD8+
C
p=0.04
p=0.0036
p=0.01
Immune phenotype
GLYCOLYSIS
UNFOLDED_PROTEIN_RESPONSE
MTORC1_SIGNALING
DNA_REPAIR
MITOTIC_SPINDLE
SPERMATOGENESIS
G2M_CHECKPOINT
E2F_TARGETS
OXIDATIVE_PHOSPHORYLATION
MYC_TARGETS_V1
MYC_TARGETS_V2
INTERFERON_ALPHA_RESPONSE
INTERFERON_GAMMA_RESPONSE
ALLOGRAFT_REJECTION
IL6_JAK_STAT3_SIGNALING
INFLAMMATORY_RESPONSE
TNFA_SIGNALING_VIA_NFKB
KRAS_SIGNALING_UP
COMPLEMENT
IL2_STAT5_SIGNALING
PROTEIN_SECRETION
PI3K_AKT_MTOR_SIGNALING
CHOLESTEROL_HOMEOSTASIS
UV_RESPONSE_UP
ESTROGEN_RESPONSE_EARLY
ESTROGEN_RESPONSE_LATE
REACTIVE_OXYGEN_SPECIES_PATHWAY
PEROXISOME
ADIPOGENESIS
FATTY_ACID_METABOLISM
HYPOXIA
ANDROGEN_RESPONSE
APOPTOSIS
P53_PATHWAY
PANCREAS_BETA_CELLS
KRAS_SIGNALING_DN
BILE_ACID_METABOLISM
XENOBIOTIC_METABOLISM
HEME_METABOLISM
EPITHELIAL_MESENCHYMAL_TRANSITION
ANGIOGENESIS
WNT_BETA_CATENIN_SIGNALING
NOTCH_SIGNALING
TGF_BETA_SIGNALING
HEDGEHOG_SIGNALING
UV_RESPONSE_DN
COAGULATION
APICAL_SURFACE
MYOGENESIS
APICAL_JUNCTION
GSVA enrichment score
Patient
HRD status
Immune phenotype
purely-inflamed
mixed-inflamed
excluded
desert
HRD status
BRCA1
BRCA2
WT
Unknown
d
M1 like
0.042
M2 like
Matrix_remodeling
Weston_VEGF targets
0.072
WNT_beta catenin signaling
0.186

**Figure S2: Excluded and desert OC exhibit higher expression of signatures related to angiogenesis and cancer progression.**

**(A)** Bar plots showing the percentage of each TILs subset depicted by mIF in the four immune phenotypes and split by tumour and stromal areas. Numbers in the upper line indicate the median $CD8^+$ T cell density. **(B)** Boxplots showing proportion of the T cell subset of interest out of total $CD8^+$ T cell profiled by mIF according to immune phenotypes and split by tumour and stromal areas; p-value assessed using unpaired, two-tailed Wilcoxon-rank test. **(C)** Heatmap of Hallmarks pathways (Table S2) showing different clustering among the four immune phenotypes. **(D)** Boxplots showing selected significant different pathways among the four immune categories (extracted from panel d); p-value assessed using unpaired, two-tailed Wilcoxon-rank test, adjusted for Bonferroni correction.

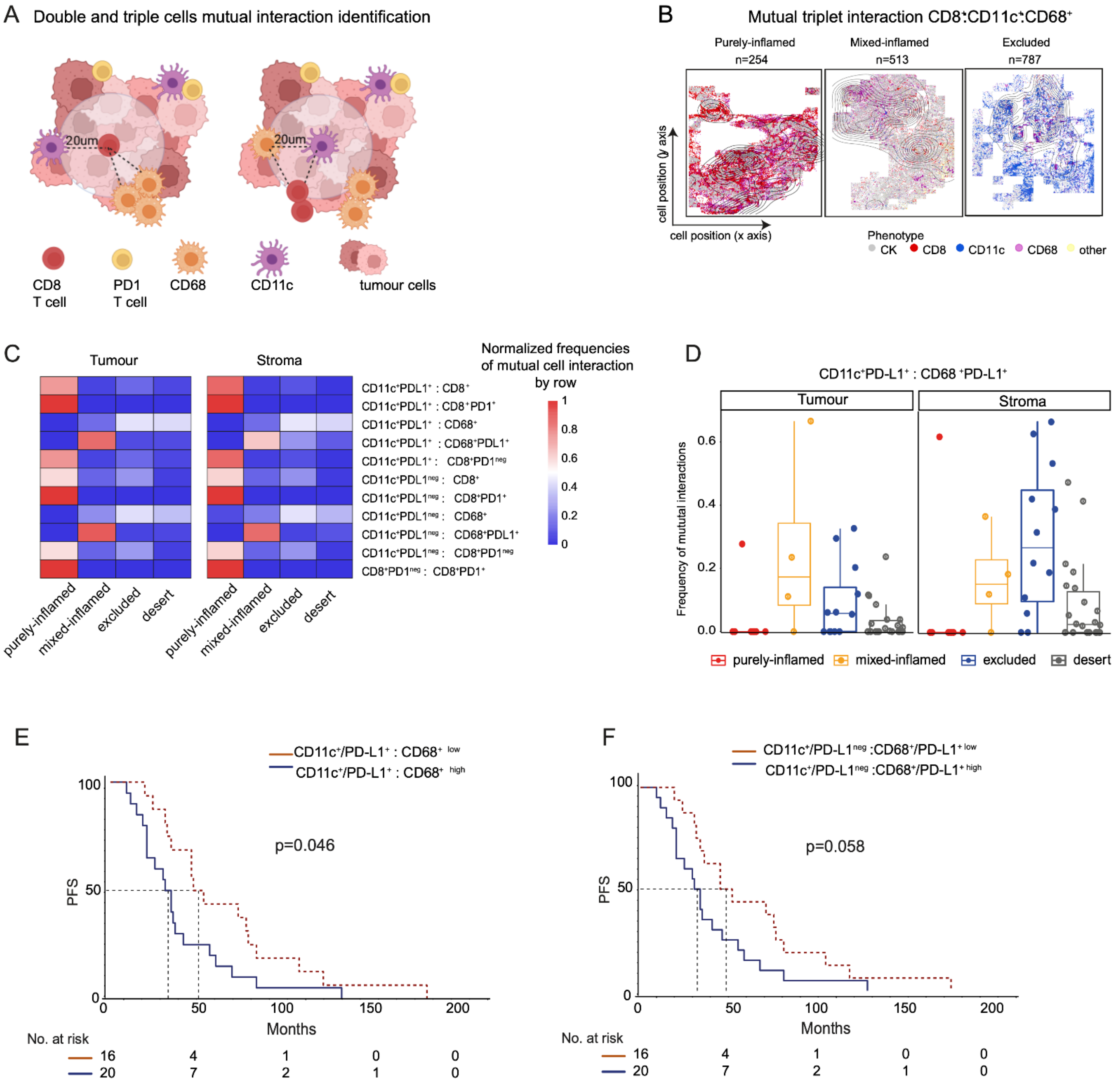
A
Double and triple cells mutual interaction identification
20um
20um
CD8
T cell
PD1
T cell
CD68
CD11c
tumour cells
B
Mutual triplet interaction CD8⁺:CD11c⁺:CD68⁺
Purely-inflamed
n=254
Mixed-inflamed
n=513
Excluded
n=787
cell position (y axis)
cell position (x axis)
Phenotype
CK
CD8
CD11c
CD68
other
C
Tumour
Stroma
Normalized frequencies
of mutual cell interaction
by row
1
0.8
0.6
0.4
0.2
0
CD11c⁺PDL1⁺ : CD8⁺
CD11c⁺PDL1⁺ : CD8⁺PD1⁺
CD11c⁺PDL1⁺ : CD68⁺
CD11c⁺PDL1⁺ : CD68⁺PDL1⁺
CD11c⁺PDL1⁺ : CD8⁺PD1ⁿᵉᵍ
CD11c⁺PDL1ⁿᵉᵍ : CD8⁺
CD11c⁺PDL1ⁿᵉᵍ : CD8⁺PD1⁺
CD11c⁺PDL1ⁿᵉᵍ : CD68⁺
CD11c⁺PDL1ⁿᵉᵍ : CD68⁺PDL1⁺
CD11c⁺PDL1ⁿᵉᵍ : CD8⁺PD1ⁿᵉᵍ
CD8⁺PD1ⁿᵉᵍ : CD8⁺PD1⁺
purely-inflamed
mixed-inflamed
excluded
desert
purely-inflamed
mixed-inflamed
excluded
desert
D
CD11c⁺PD-L1⁺ : CD68⁺PD-L1⁺
Tumour
Stroma
Frequency of mutual interactions
0.6
0.4
0.2
0.0
purely-inflamed
mixed-inflamed
excluded
desert
E
CD11c⁺/PD-L1⁺ : CD68⁺ low
CD11c⁺/PD-L1⁺ : CD68⁺ high
p=0.046
PFS
100
50
0
0
50
100
150
200
Months
No. at risk
16 4 1 0 0
20 7 2 1 0
F
CD11c⁺/PD-L1ⁿᵉᵍ :CD68⁺/PD-L1⁺ low
CD11c⁺/PD-L1ⁿᵉᵍ :CD68⁺/PD-L1⁺ high
p=0.058
PFS
100
50
0
0
50
100
150
200
Months
No. at risk
16 4 1 0 0
20 7 2 1 0

**Figure S3: Mixed-inflamed and excluded tumours exhibit higher homotypic myeloid interactions at baseline affecting overall survival.**

**(A)** Cartoon displaying double and triple mutual interaction within the 20um radius as described in Methods. **(B)** Representative images of digital tissue reconstruction showing the density of the frequency of triple mutual interaction $CD8^{+}$:$CD68^{+}$:$CD11c^{+}$ according to immune phenotypes. **(C)** Heatmaps showing the differences in the frequency of mutual interaction between cell types (including PD1+ and PD-L1+ subpopulations) at a 20um neighboring radii in the tumour and stroma compartment. Immune cell population interaction of interest in lines and immune categories as columns. Color-code scale bar showing the normalized frequency by each row. **(D)** Boxplots displaying frequency of mutual interaction between the indicated cell types according to immune phenotypes and split by tumour or stromal; p-value assessed using unpaired, two tailed Wilcoxon-rank test. **(E-F**) Kaplan-Meier curve of OS in the UPENN cohort according to homotypic niches enrichment, $CD11c^{+}$PD-$L1^{+}$ :$CD68^{+}$ and $CD11c^{+}$ PD-$L1^{NEG}$ :$CD68^{+}$PD-$L1^{+}$ respectively; p value assessed using the Log-rank test.

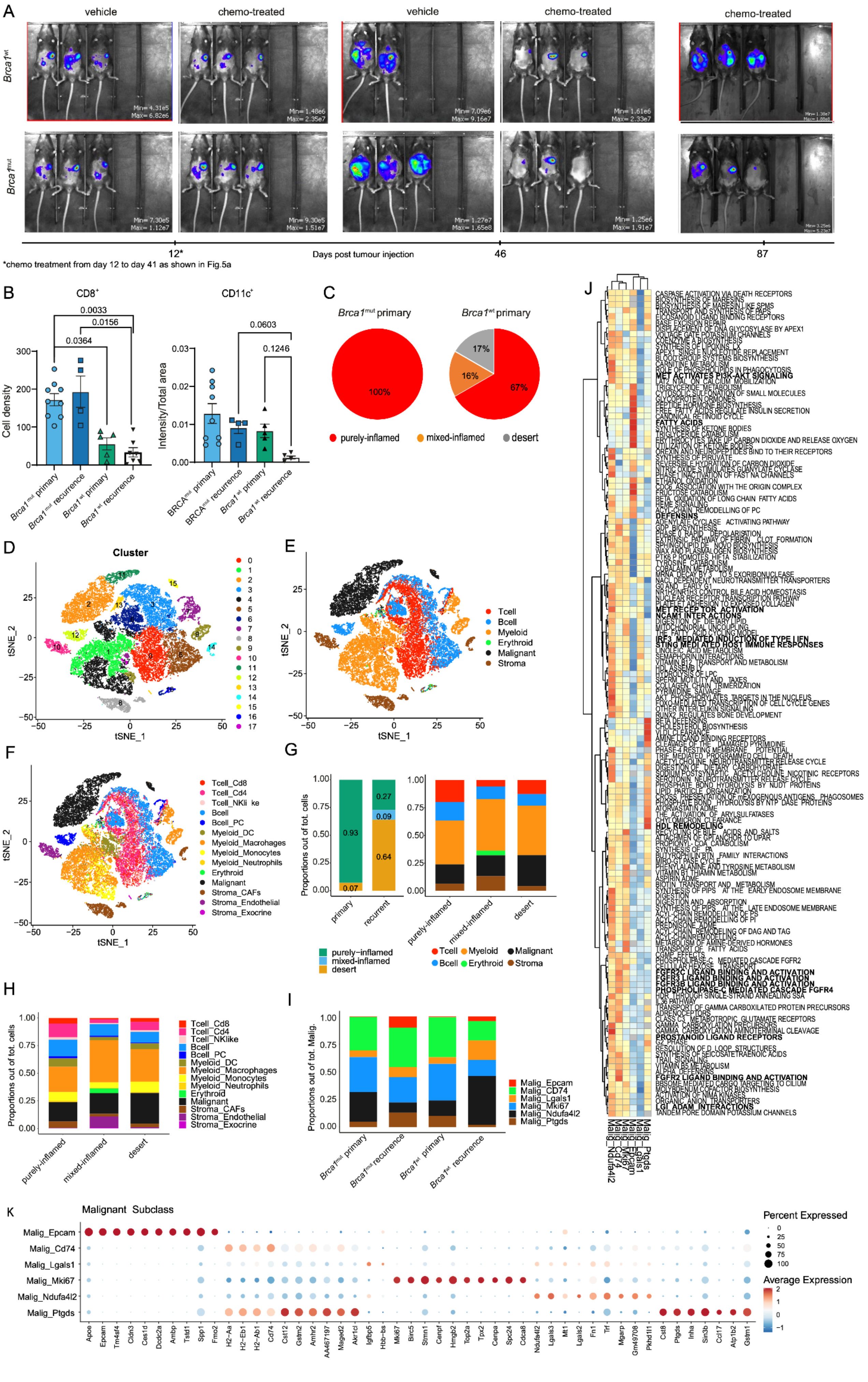

A
vehicle
chemo-treated
vehicle
chemo-treated
chemo-treated
Brca1wt
Brca1mut
12*
Days post tumour injection
46
87
*chemo treatment from day 12 to day 41 as shown in Fig.5a
B
CD8+
CD11c+
Cell density
Intensity/Total area
0.0033
0.0156
0.0364
0.0603
0.1246
Brca1mut primary
Brca1mut recurrence
Brca1wt primary
Brca1wt recurrence
C
Brca1mut primary
Brca1wt primary
100%
17%
16%
67%
purely-inflamed
mixed-inflamed
desert
D
Cluster
tSNE_1
tSNE_2
E
Tcell
Bcell
Myeloid
Erythroid
Malignant
Stroma
F
Tcell_Cd8
Tcell_Cd4
Tcell_NKlike
Bcell
Bcell_PC
Myeloid_DC
Myeloid_Macrohages
Myeloid_Monocytes
Myeloid_Neutrophils
Erythroid
Malignant
Stroma_CAFs
Stroma_Endothelial
Stroma_Exocrine
G
Proportions out of tot. cells
primary
recurrent
0.93
0.07
0.27
0.09
0.64
purely-inflamed
mixed-inflamed
desert
H
Proportions out of tot. cells
I
Proportions out of tot. Malig.
Malig_Epcam
Malig_CD74
Malig_Lgals1
Malig_Mki67
Malig_Ndufa4l2
Malig_Ptgds
J
K
Malignant Subclass
Percent Expressed
Average Expression
Apoe
Epcam
Tm4sf4
Cldn3
Ces1d
Dcdc2a
Ambp
Tstd1
Spp1
Fmo2
H2-Aa
H2-Eb1
H2-Ab1
Cd74
Cst12
Gstm2
Amhr2
AA467197
Maged2
Akr1cl
Igfbp5
Hbb-bs
Mki67
Birc5
Stmn1
Cenpf
Hmgb2
Top2a
Tpx2
Cenpa
Spc24
Cdca8
Ndufa4l2
Lgals3
Mt1
Lgals2
Fn1
Trf
Mgarp
Gm49708
Pkhd1l1
Cst8
Ptgds
Inha
Sin3b
Ccl17
Atp1b2
Gstm1

**Figure S4: *Brca1$^{mut}$* and *Brca1$^{wt}$* tumours show divergent tumoral and TME states upon chemotherapy.**

**(A)** Example of mice imaging follow-up acquired by luciferase signaling, referring to experiment in Figure 5A. **(B)** Boxplots showing CD8+ and CD11c$^{+}$ cell density assessed by IHC between *Brca1$^{mut}$* and *Brca1$^{wt}$* tumours at baseline and at recurrence; p-value assessed using unpaired, two-tailed Wilcoxon-rank test. **(C)** Pie-charts displaying the different percentage of immune phenotypes at baseline between *Brca1$^{mut}$* and *Brca1$^{wt}$* tumours. **(D)** t-SNE map of single-cell data reveals 17 major cell clusters. e-f t-SNE map of single-cell data showing major cell types and subtypes, respectively. **(G-H)** Bar plots displaying respectively the proportion of cell types and subtypes (out of total cells assigned) according to immune-phenotypes. **(I)** Bar plots displaying the proportion of malignant subtypes between Brca1mut and Brca1wt tumours at baseline and at recurrence. **(J)** Heatmap showing differential gene expression pathways among the six different malignant subclasses. **(K)** Bubble plot of gene expression of the top 10 differential genes for each identified malignant subpopulation.

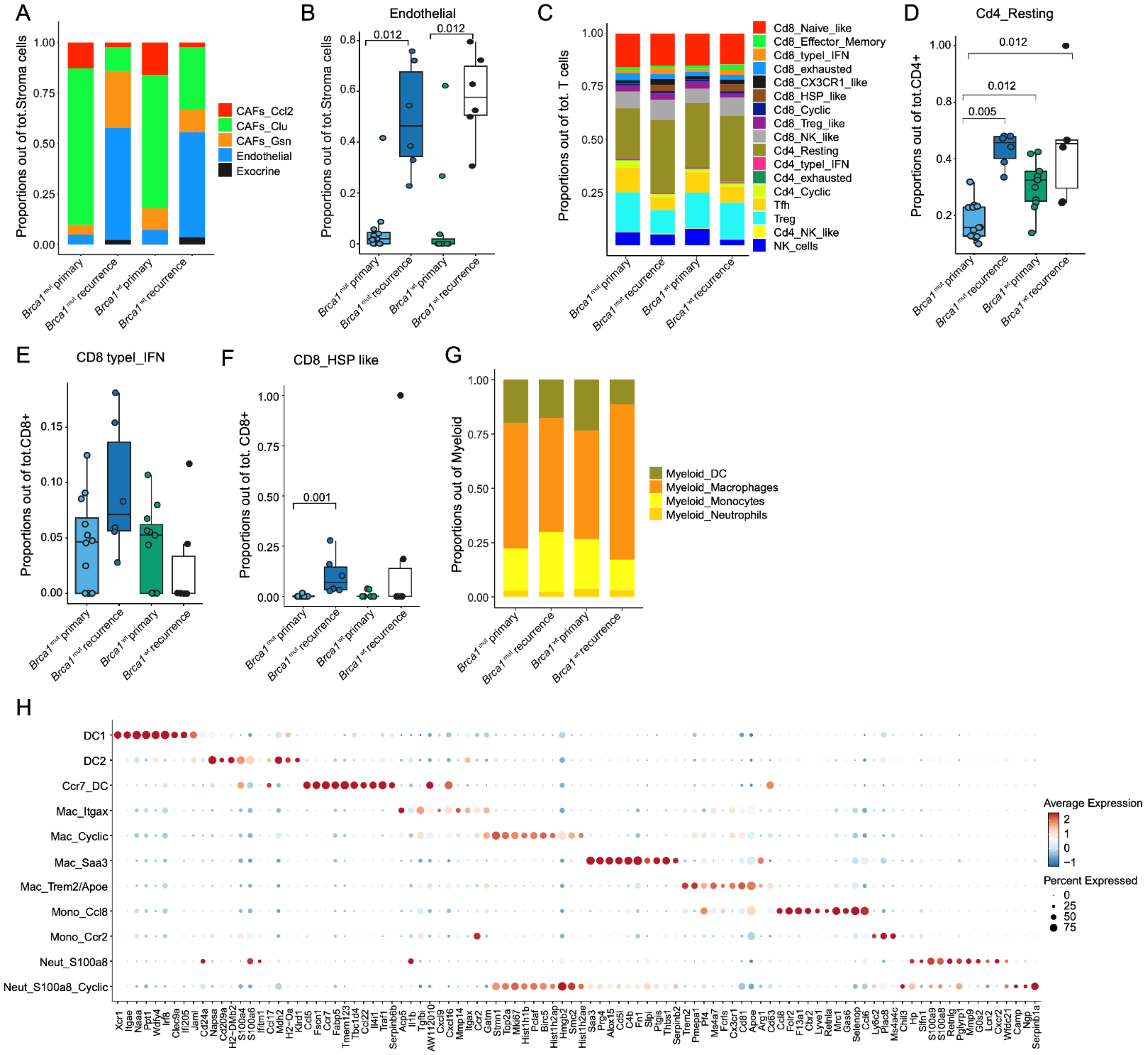

A
Proportions out of tot.Stroma cells
CAFs_Ccl2
CAFs_Clu
CAFs_Gsn
Endothelial
Exocrine
Brca1 mut primary
Brca1 mut recurrence
Brca1 wt primary
Brca1 wt recurrence
B
Endothelial
0.012
0.012
Proportions out of tot.Stroma cells
C
Proportions out of tot. T cells
Cd8_Naive_like
Cd8_Effector_Memory
Cd8_typel_IFN
Cd8_exhausted
Cd8_CX3CR1_like
Cd8_HSP_like
Cd8_Cyclic
Cd8_Treg_like
Cd8_NK_like
Cd4_Resting
Cd4_typel_IFN
Cd4_exhausted
Cd4_Cyclic
Tfh
Treg
Cd4_NK_like
NK_cells
D
Cd4_Resting
0.012
0.012
0.005
Proportions out of tot.CD4+
E
CD8 typel_IFN
Proportions out of tot.CD8+
F
CD8_HSP like
0.001
Proportions out of tot. CD8+
G
Proportions out of Myeloid
Myeloid_DC
Myeloid_Macrophages
Myeloid_Monocytes
Myeloid_Neutrophils
H
DC1
DC2
Ccr7_DC
Mac_Itgax
Mac_Cyclic
Mac_Saa3
Mac_Trem2/Apoe
Mono_Ccl8
Mono_Ccr2
Neut_S100a8
Neut_S100a8_Cyclic
Average Expression
Percent Expressed

**Figure S5: *Brca*$^{mut}$ inflamed tumours retain TILs and DC interactions upon CXT.**

**(A)** Bar plots displaying the proportion of stromal cell type and subtypes (out of total cells) among *Brca1*$^{mut}$ and *Brca1*$^{wt}$ tumours at baseline and at recurrence. **(B)** Boxplots displaying the proportion of endothelial cell between *Brca1*$^{mut}$ and *Brca1*$^{wt}$ at baseline and at recurrence; p-value assessed using unpaired, two-tailed Wilcoxon-rank test, adjusted for Bonferroni correction. **(C)** Bar plots displaying the proportion of T cells subclasses (out of total T cells) among *Brca1*$^{mut}$ and *Brca1*$^{wt}$ tumours at baseline and at recurrence. **(D-F)** Boxplots displaying the proportion of the indicated T cell subclass between *Brca1*$^{mut}$ and *Brca1*$^{wt}$ at baseline and at recurrence; p-value assessed using unpaired, two-tailed Wilcoxon-rank test, adjusted for Bonferroni correction. **(G)** Bar plots displaying the proportion of myeloid cell subtypes (out of total myeloid cells) among *Brca1*$^{mut}$ and *Brca1*$^{wt}$ at baseline and at recurrence. **(H)** Bubble plot of gene expression of the top 10 differential genes for each identified myeloid subclass.

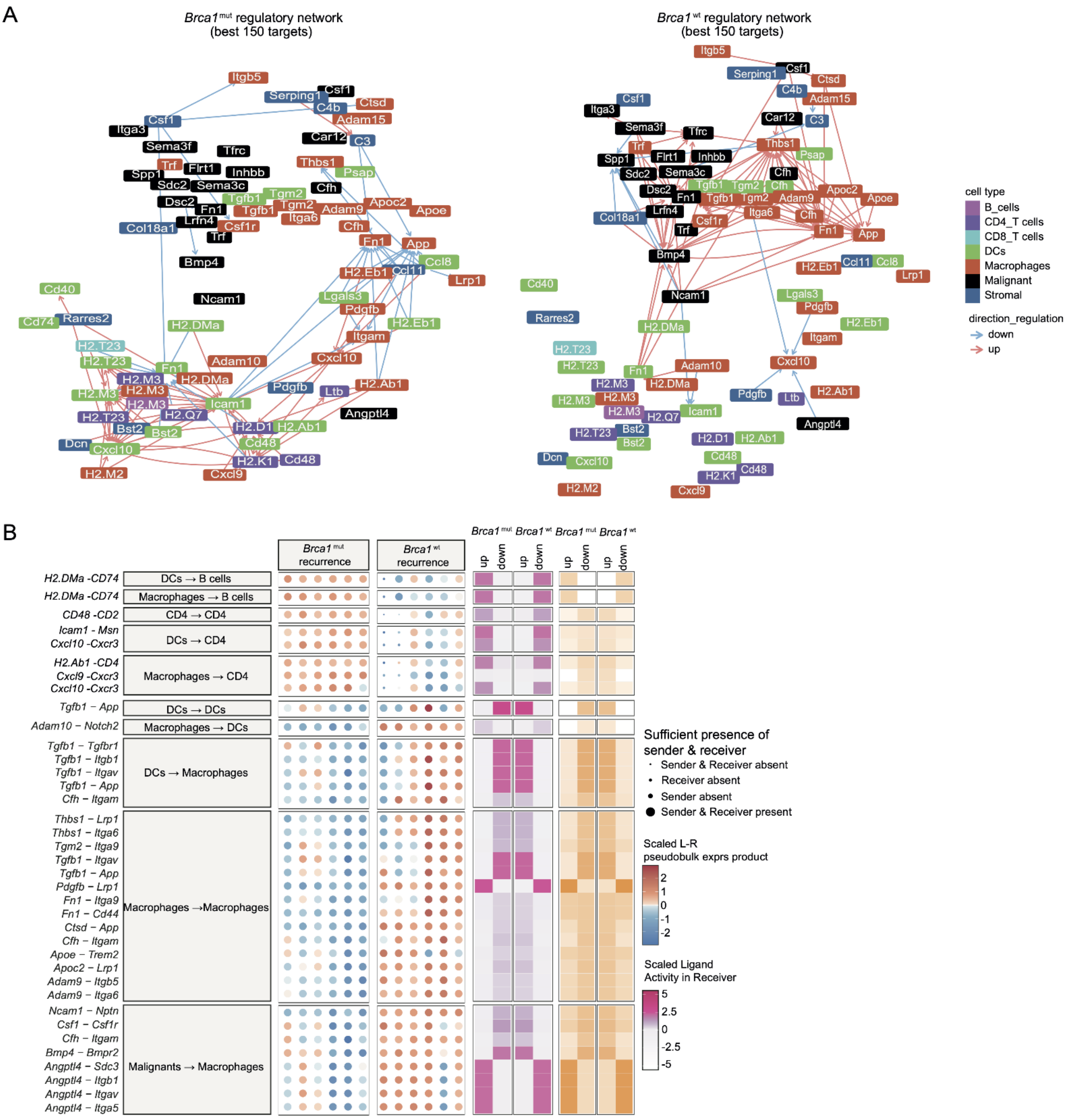
A
Brca1 mut regulatory network
(best 150 targets)
Brca1 wt regulatory network
(best 150 targets)
cell type
B_cells
CD4_T cells
CD8_T cells
DCs
Macrophages
Malignant
Stromal
direction_regulation
down
up
B
Brca1 mut recurrence
Brca1 wt recurrence
Brca1 mut
Brca1 wt
Brca1 mut
Brca1 wt
up
down
up
down
up
down
up
down
H2.DMa -CD74
DCs → B cells
H2.DMa -CD74
Macrophages → B cells
CD48 -CD2
CD4 → CD4
Icam1 - Msn
Cxcl10 -Cxcr3
DCs → CD4
H2.Ab1 -CD4
Cxcl9 -Cxcr3
Cxcl10 -Cxcr3
Macrophages → CD4
Tgfb1 – App
DCs → DCs
Adam10 – Notch2
Macrophages → DCs
Tgfb1 – Tgfbr1
Tgfb1 – Itgb1
Tgfb1 – Itgav
Tgfb1 – App
Cfh – Itgam
DCs → Macrophages
Thbs1 – Lrp1
Thbs1 – Itga6
Tgm2 – Itga9
Tgfb1 – Itgav
Tgfb1 – App
Pdgfb – Lrp1
Fn1 – Itga9
Fn1 – Cd44
Ctsd – App
Cfh – Itgam
Apoe – Trem2
Apoc2 – Lrp1
Adam9 – Itgb5
Adam9 – Itga6
Macrophages →Macrophages
Ncam1 – Nptn
Csf1 – Csf1r
Cfh – Itgam
Bmp4 – Bmpr2
Angptl4 – Sdc3
Angptl4 – Itgb1
Angptl4 – Itgav
Angptl4 – Itga5
Malignants → Macrophages
Sufficient presence of
sender & receiver
Sender & Receiver absent
Receiver absent
Sender absent
Sender & Receiver present
Scaled L-R
pseudobulk exprs product
2
1
0
-1
-2
Scaled Ligand
Activity in Receiver
5
2.5
0
-2.5
-5

**Figure S6: Tgfb1, Fn1, App and Thbs1 signaling drive immunosuppressive TAM networks in recurrent *Brca1$^{wt}$* tumours.**

**(A)** Differential regulatory network (best 150 targets) predicted by MultiNicheNet analysis in the *Brca1$^{mut}$* and *Brca1$^{wt}$* mouse models at recurrence. **(B)** Bubble-plot of interactome analysis as by prioritized top 50 interactions in *Brca1$^{mut}$* and *Brca1$^{wt}$* tumours (here only the top 5 cell type interaction showed); detailed ligand interactions are listed in the left column.

#### Supplementary Tables

- Table S1. IMCOL and UPENN primary-recurrent cohorts' patients and samples characteristics.
- Table S2. HiTide-UPENN cohort' patients and samples characteristics, including gene signature analysis (referring to Figures 2D and S2C).
- Table S3. MultiNicheNet analysis from single-cell RNAsequencing on human samples (HiTide-UPENN cohort, n=19, referring to Figure 3J).
- Table S4. Detailed single-cellRNA sequencing analysis subclasses gene profiling on mouse models.
- Table S5. MultiNicheNet analysis on mouse samples (referring to Figures 6E and S6A-B).
